# Targeting poorly differentiated thyroid cancer with bispecific antibodies and CAR T cells from the human anti-TSHR autoantibody repertoire

**DOI:** 10.64898/2026.10.19.760040

**Authors:** Claudia Fischer, Anne Eugster, Brenda Besemer, Mélanie Buchi, Marcus Bauer, Christoph Schultheiß, Elisa D’Arcangelo, Filip Filipsky, Claudia Wickenhauser, Andreas Wilfer, Kerstin Lorenz, Jens Jakscha, Laurent Muller, Roswitha Köberle-Wührer, Benjamin Kasenda, Christof Rottenburger, Rauf Latif, Terry F. Davies, Heinz Läubli, Matthias Peipp, Mascha Binder

**Affiliations:** Division of Medical Oncology, University Hospital Basel, Basel, Switzerland; Laboratory of Translational Immuno-Oncology, Department of Biomedicine, University of Basel and University Hospital of Basel, Basel, Switzerland; Laboratory of Cancer Immunology, Department of Biomedicine, University of Basel and University Hospital of Basel, Basel, Switzerland; Institute of Pathology, University Hospital Halle (Saale), Martin-Luther-University Halle-Wittenberg, Halle (Saale), Germany; Institute of Human Biology (IHB), Roche Pharma Research and Early Development, Basel, Switzerland; Laboratory of Cancer Immunotherapy, Department of Biomedicine, University of Basel and University Hospital of Basel, Basel, Switzerland; Department of Visceral, Vascular and Endocrine Surgery, Medical Faculty, Martin-Luther-University Halle-Wittenberg, Halle (Saale), Germany; Department of Otolaryngology, Head and Neck Surgery, University Hospital Basel, Basel, Switzerland; Department of Visceral Surgery, Claraspital Basel, Clarunis, Basel, Switzerland; Division of Nuclear Medicine, University Hospital Basel, Basel, Switzerland; Thyroid Research Unit, Icahn School of Medicine at Mount Sinai, New York, NY, USA; James J. Peters VA Medical Center, New York, NY, USA; Innovation Focus Cell Therapies, University Hospital Basel, Basel, Switzerland; Division of Antibody-Based Immunotherapy, Department of Internal Medicine II (Hematology/Oncology), University Hospital Schleswig-Holstein Campus Kiel, Kiel, Germany; Collaborative Research Institute Intelligent Oncology (CRIION), Freiburg, Germany

## Abstract

**Background:** Poorly differentiated thyroid carcinoma (PDTC) has no effective systemic treatment option once radioactive iodine and tyrosine kinase inhibitors fail. The thyroid-stimulating hormone receptor (TSHR) is highly tissue-restricted and the anti-TSHR autoantibody repertoire of thyroid autoimmunity offers sequence-defined, fully human binders. We asked whether this repertoire can be converted into an off-the-shelf T cell-redirecting therapy for PDTC.

**Methods:** TSHR expression was profiled on a 151-case thyroid cancer tissue microarray (TMA), 50 healthy controls and on fresh tumor material. Eight anti-TSHR clones, four of human autoimmune origin (M22, K1-18, K1-70, 5C9) and four murine (IRI-SAb2, IRI-SAb3, 9D33, 1H7), were reformatted as CD3-engaging bispecific antibodies (bsAbs; anti-TSHR Fab × anti-CD3 scFv) and, for leading clones, as chimeric antigen receptor (CAR) T cells. Activity was compared *in vitro*, across graded and mixed antigen densities and in a CAL-62-Luc-TSHR xenograft model, using healthy-donor (HD) or thyroid cancer-patient T cells.

**Results:** TSHR was retained on a substantial fraction of PDTC, including metastatic lesions. Both formats mediated potent, antigen-selective killing and the leading bsAbs suppressed xenograft growth *in vivo*. For CAR T cells, scFv heavy/light-chain orientation was decisive. Anti-TSHR bsAbs cleared mixed cultures far beyond the antigen-positive fraction matching mosunetuzumab and indicating a modality-intrinsic bystander effect. Patient-derived T cells obtained on or after oncological treatment retained full effector function.

**Conclusions:** The anti-TSHR autoantibody repertoire provides a promising source of binders for off-the-shelf bispecific antibodies targeting TSHR-positive thyroid cancer, while CAR T cells offer an autologous alternative.

## INTRODUCTION

The incidence of thyroid cancer has been rising globally over the past decade, with most cases belonging to differentiated thyroid carcinomas (DTC), as follicular (FTC) or papillary (PTC) (1). The majority of patients with DTC achieve durable disease control after surgery and radioactive iodine (RAI) therapy (2). In contrast, poorly differentiated thyroid carcinoma (PDTC) and anaplastic thyroid carcinoma (ATC) carry a markedly worse prognosis (3). PDTC, while less devastating than ATC, frequently becomes refractory to RAI and progresses despite tyrosine kinase inhibitor (TKI) therapy (3). Clinically, the unmet need is greatest in these advanced and metastatic settings where conventional cytotoxic chemotherapy is of limited benefit and BRAF-, RET- or NTRK-directed agents are only applicable in a minority of patients (4–6).

The thyroid-stimulating hormone receptor (TSHR) is a G-protein-coupled receptor whose physiological expression is mostly confined to thyroid follicular cells (7). TSHR expression in DTC is reliable but typically lost upon full anaplastic dedifferentiation (7). Expression in PDTC is less well characterized but has been reported to be reduced compared to normal thyroid tissue (7, 8). The tissue specificity of TSHR makes it an attractive candidate antigen for engineered immunotherapeutic targeting (9).

Targeting moieties of suitable specificity and affinity already exist. In autoimmune thyroid disease, B cell-derived autoantibodies recognize multiple thyroid antigens including TSHR (10–12). Anti-TSHR antibodies in Graves’ disease and primary hypothyroidism span a remarkable functional spectrum from thyroid-stimulating (e.g. M22, K1-18) to blocking (e.g. K1-70, 5C9, 9D33) and their interactions with TSHR have been mapped at high resolution (13–15). Two clinically validated antibody-based effector platforms are particularly suited to exploit such binders. CD3-engaging bispecific antibodies (bsAbs) physically bridge tumor cells and T cells and have transformed the treatment of B cell lymphoma, multiple myeloma and small-cell lung cancer (16–18), and the engineering principles for such formats are well established (19). CAR T cells, in turn, are now standard of care in B cell and plasma-cell malignancies and are being intensely explored in solid tumors and autoimmune disease (20, 21). TSHR-directed CAR T cells have been reported for differentiated thyroid cancer using either an antibody-derived scFv or the TSH ligand itself as the targeting moiety (22–25). These studies, however, addressed differentiated thyroid cancers and did not compare bsAb and CAR formats built from the same variable-region repertoire, and did not use T cells from the intended patient population (24). No CD3-engaging bispecific antibody against TSHR has been described in literature.

Here, we set out to lay the preclinical foundation for TSHR-directed immunotherapy in PDTC. Specifically, we re-examined TSHR expression across the full spectrum of thyroid cancer histologies on a 151-case tissue microarray (TMA) and on fresh tumor material. We curated eight anti-TSHR antibody clones spanning the stimulating and blocking pharmacological classes, four of human autoimmune origin, and took all eight variable-region sequences forward to generate bsAbs, and the validated subset forward to generate CAR T cells. Comparing their target engagement, cytolytic activity and effector signatures across two tumor cell systems helped with narrowing down our selection to the two most promising candidates. These were further investigated against graded TSHR expression levels and in mixed populations of TSHR-positive and TSHR-negative target cells modeling intratumoral antigen heterogeneity. The leading bsAb constructs were additionally investigated in a CAL-62-Luc-TSHR xenograft model. Finally, T cells from patients with thyroid cancer were found to be capable of mediating the cell death induced by anti-TSHR bsAb or of being reprogrammed into anti-TSHR CAR-T cells.

## METHODS

### Patients and healthy donors

A retrospective, anonymized thyroid cancer cohort was assembled with the Institute of Pathology and the Department of Visceral, Vascular and Endocrine Surgery at University Hospital Halle as described (26). Patients diagnosed with PTC (n = 84), FTC (n = 29), PDTC (n = 18) and ATC (n = 20) and healthy, non-neoplastic thyroid tissue from 50 control individuals as reference were included in this study. Fresh tumor tissue and patient-derived T cells were obtained from the Department of Otolaryngology, Head and Neck Surgery, University Hospital Basel, and T cells from age-matched healthy donors (HD) from the Blood Donation Center Both Basel (Blutspendezentrum beider Basel). All donors gave written informed consent in accordance with the Declaration of Helsinki. The study was approved by the Ethics Committee of the Medical Faculty Halle (2017–81) and by EKNZ, Basel. Pan T cells were isolated by Ficoll gradient centrifugation and magnetic bead sorting (Miltenyi).

### TSHR immunohistochemistry on FFPE tissue

A tissue microarray was prepared from 151 formalin-fixed, paraffin-embedded (FFPE) thyroid cancer and 50 non-neoplastic thyroid samples. Immunohistochemistry on 3 µm sections was performed on a fully automated Leica Bond stainer using two anti-TSHR antibodies: EPR19751 (Abcam, RRID:AB_3096105, Cat# ab218108, 1:1000), shown for the thyroid cancer tissue microarray and the non-neoplastic control tissue in Figure 1A, B, and 4C1 (Santa Cruz, RRID:AB_628357, Cat# sc-32262, 1:100), shown for the same tissue microarray and for the engineered CAL-62 cell lines in Supplementary Figure 9C. Staining was scored by two pathologists blinded to clinical data and expressed as an H-score (percentage of positive tumor cells × intensity 0–3), classified as absent (0), low (1–100), intermediate (101–200) or strong (201–300). Staining conditions are detailed in Supplementary Methods.

**Figure 1.**
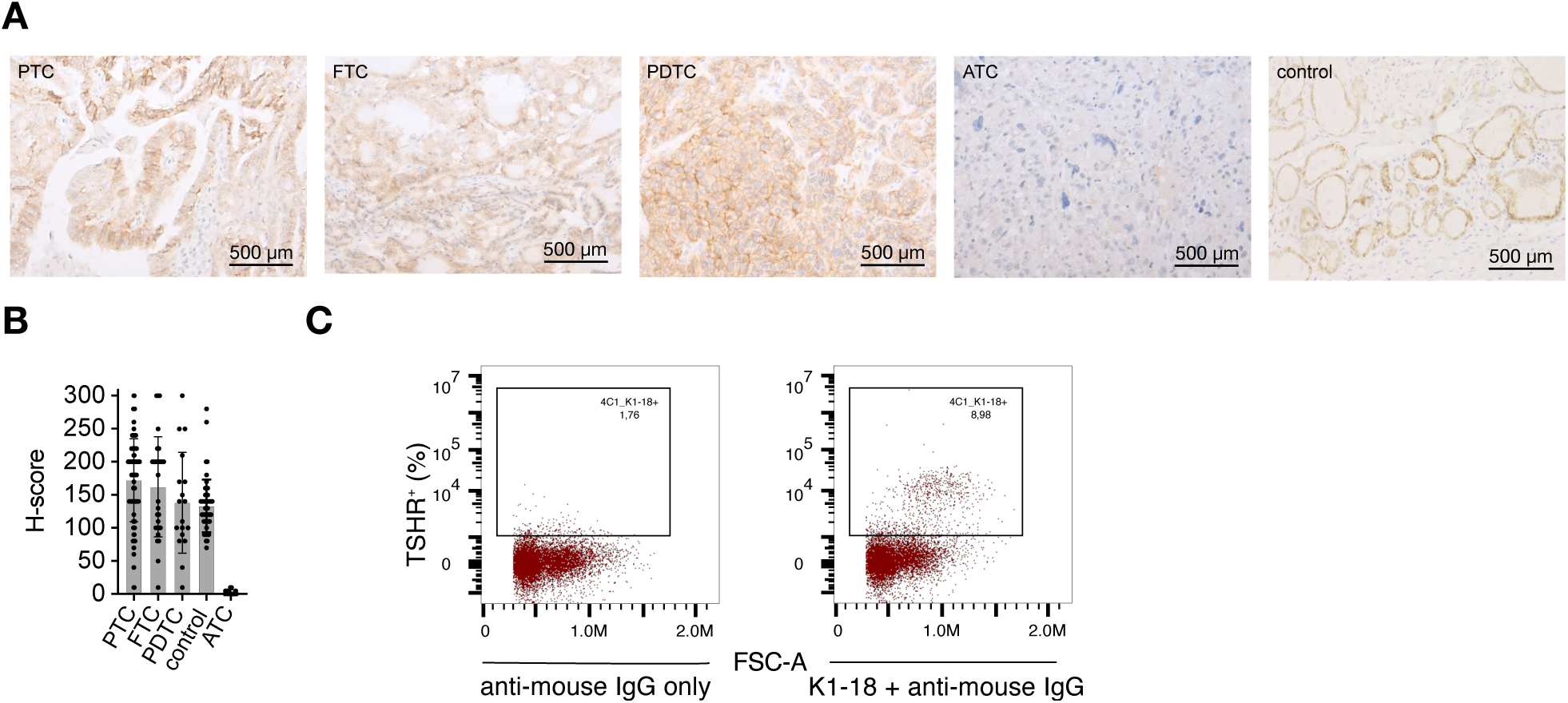
TSHR expression in primary thyroid cancer tissue. **A** Representative images of TSHR immunohistochemistry (anti-TSHR antibody clone EPR19751) for each thyroid cancer subgroup and for the control, i.e. normal, non-neoplastic thyroid tissue. **B** H-score-based quantification of TSHR immunohistochemistry (clone EPR19751) on a tissue microarray of thyroid cancer cases (PTC, n = 84; FTC, n = 29; PDTC, n = 18; ATC, n = 20). Error bars show mean ± SD with individual cases overlaid. **C** Flow-cytometric detection of surface TSHR on dissociated fresh tumor material, stained with the leucine-rich-repeat-directed antibody K1-18 followed by an APC-conjugated anti-mouse IgG secondary antibody. Staining with the secondary antibody alone is shown as a control. PTC: papillary thyroid carcinoma; FTC: follicular thyroid carcinoma; PDTC: poorly differentiated thyroid carcinoma; ATC: anaplastic thyroid carcinoma.

### TSHR expression detection on fresh tissue

Fresh tumor tissue was mechanically dissociated and enzymatically digested for 1 h to overnight at 37 °C, filtered through a 100 µm strainer and cryopreserved. Surface TSHR was detected by flow cytometry using the chimeric anti-TSHR antibody K1-18 (AntibodySystem, Cat# VHD18101) and an APC-conjugated anti-mouse IgG secondary antibody (Jackson ImmunoResearch, RRID:AB_2338648, Cat# 115-136-071). The digestion mix is specified in Supplementary Methods.

### Cell lines and target cell engineering

The ATC cell lines CAL-62 and C643 (DSMZ, CVCL_1112, CVCL_5969) were lentivirally transduced with constructs encoding firefly luciferase (luc2) and GFP, human MS4A1 (CD20) or full-length TSHR under the SFFV promoter. Graded TSHR expression was obtained by transducing CAL-62 cells with constructs carrying an IRES sequence with mutated AUG-12 (IRESv3) or AUG-11 (IRESv7) preceding the TSHR cDNA (27), or with the TSHR cassette under the control of the inducible TRE3GS promoter using the TetOn 3G system (28). Resulting cells are referred to as CAL-62-Luc-IRESv3-TSHR, CAL-62-Luc-IRESv7 and CAL-62-Luc-tetTSHR cells. TSHR expression was checked regularly by flow cytometry staining using either the anti-TSHR 4C1 as described above or the anti-TSHR M22 (MedChemExpress, Cat# HY-P991202) and an APC-Cy7-conjugated anti-human IgG Fc secondary antibody (BioLegend, Cat# 410732). Lentiviral production and cell transduction is detailed in Supplementary Methods.

### Bispecific antibody design

Variable- and joining-region sequences (IGHV/J, IGKV/J, IGLV/J) and CDR3 regions of the human anti-TSHR autoantibodies M22, K1-18, K1-70 and 5C9 and of the murine antibodies IRI-SAb2, IRI-SAb3, 9D33 and 1H7 were taken from the literature and the corresponding patent disclosures (29–32) and assembled as Fab-format TSHR-targeting arms. Heavy-chain variable regions (VH) were fused to an IgG1 Fc carrying the knob mutations S354C/T366W and the Fc-silencing L234A/L235A substitutions (33, 34). Light-chain variable regions (VL) were fused to their full constant region. The CD3 arm comprised the UCHT1 variable regions as a single-chain variable fragment (scFv) (35) joined through a truncated IgG1 hinge to an Fc carrying the hole mutations Y349C/T366S/L368A/Y407V. All constructs were fused to a 5’ IL-2 leader sequence and cloned into pcDNA3(+). Linker and cloning details are given in Supplementary Methods.

### Bispecific antibody production and purification

Bispecific antibodies were produced in FreeStyle™ CHO-S cells (Thermo Fisher) by electroporation, harvested after at most 14 days, and purified by affinity capture on a HiTrap™ Fibro PrismA column followed by size-exclusion chromatography on a Superdex 200 Increase 10/300 GL column (both Cytiva). Monomer fractions were pooled in PBS and purity was verified by SDS-PAGE with Coomassie staining (Bio-Rad). Culture, feeding and harvest conditions are detailed in Supplementary Methods.

### CAR constructs and lentiviral production

Second-generation anti-TSHR CAR constructs (backbone pJ2459) carry the 4-1BB and CD3ζ signaling domains linked through a truncated IgG4 hinge (ESKYGPPCPPCP) to the scFv of the respective anti-TSHR antibody, in both VH–VL (HL) and VL–VH (LH) orientation, under the control of the EF-1α core promoter and fused to a GM-CSF signal peptide (36, 37). Third-generation lentiviral particles were produced in Lenti-X 293T cells, concentrated by PEG-8000 precipitation and titrated by qPCR. Packaging plasmids and media are listed in Supplementary Methods.

### CAR T cell and CAR-Jurkat cell generation

Pan T cells isolated from HD or patient peripheral blood mononuclear cells were activated with CD3/CD28 Dynabeads and transduced with anti-TSHR CAR lentivirus 24 h later. Beads were removed after 48 h and cells were expanded for 10–14 days in G-Rex plates. Transduction efficiency was determined by flow cytometry using an anti-G4S-linker antibody. Activation markers (CD69, CD25, LAG-3, PD-1, TIM-3) were assessed in parallel on DAPI⁻ CD4⁺ and CD8⁺ CAR T cells. CAR Jurkat-NFAT-Luc lines were generated in the same way but without bead stimulation. Antibody clones and media are listed in Supplementary Methods.

### Jurkat-NFAT-Luc cell assays

Jurkat-NFAT-Luc reporter cells express luciferase upon CD3 and TCR pathway activation (38). Target cells (1.5 × 10⁴ CAL-62-TSHR or CAL-62-CD20) were seeded overnight, then 10 nM anti-TSHR bsAb or mosunetuzumab together with Jurkat-NFAT-Luc cells or anti-TSHR CAR Jurkat-NFAT-Luc cells, were added at an effector-to-target (E:T) ratio of 5:1 for 24 h. Monocultures and ImmunoCult-stimulated reporter cells served as controls. Bioluminescence was read on a Synergy H1 plate reader (BioTek) after addition of Bio-Glo™ reagent (Promega).

### *In vitro* cytotoxicity assays

Luciferase- and GFP-expressing target cells (1.5 × 10⁴) were seeded in 96-well plates overnight, before bispecific antibodies and effector T cells from healthy donors or thyroid cancer patients were added at an effector-to-target (E:T) ratio of 5:1 unless stated otherwise and co-cultured for 48 h. CAR T cell co-cultures used 5 × 10³ target cells per well and were analyzed after 24 h. CAR T cell amounts were adjusted for transduction efficiency. Antibody concentrations are given in the figure legends. Maximal lysis was defined by 2% Triton™ X-100 (TritonX, Sigma-Aldrich) and baseline by target cells alone. CD20-positive targets with mosunetuzumab served as a control. For TetOn experiments, target cells were pre-incubated with 0, 9, 13 or 18 ng/mL doxycycline (Merck) for 72 h and TSHR surface expression was verified by flow cytometry on days 0 and 3. Bystander killing was assessed by co-culturing defined ratios of CAL-62-Luc and CAL-62-Luc-TSHR cells. Target cell death was quantified by luciferase assay (Bio-Glo™, Promega), lactate dehydrogenase (LDH) release (Cytotoxicity Detection Kit, Roche), and IFN-γ in the supernatant by a no-wash immunoassay (Promega). Target cell proliferation was monitored by live-cell imaging (Incucyte S3, Sartorius). T cell activation was analyzed by flow cytometry for CD4, CD8, CD69, CD25, LAG-3, PD-1 and TIM-3 on DAPI⁻ GFP⁻ cells. Formulas for specific killing and LDH-based cytotoxicity are given in Supplementary Methods.

### CHO-TSHR-Luc studies

To test whether the bispecific antibodies that monovalently target the TSHR retain the stimulatory or blocking properties of their parental antibodies, CHO-TSHR-Luc reporter cells were used as described (39). Cells (1.5 × 10⁴) were seeded overnight. For stimulation, antibodies were added at varying concentrations for 4 h, with bovine thyroid-stimulating hormone (bTSH) and the stimulating antibodies M22 and K1-18 as controls. For blocking, cells were pre-incubated with antibody for 1 h before a fixed concentration of bTSH or M22 was added for a further 4 h. Bioluminescence was read after addition of Bio-Glo™ reagent and normalized to the signal from cells receiving neither bTSH nor antibody.

### *In vivo* xenograft model

Studies with mice were performed in accordance with the Swiss animal welfare regulations and approved by the Cantonal Veterinary Office of Basel-Stadt, Switzerland (licence: 3036) following the ARRIVE 2.0 guidelines, closer described in Supplementary Methods. Twenty-six 7–9-week-old NSG (NOD/ SCID/IL2Rγnull) mice were injected subcutaneously into the right flank with 5 × 10⁶ CAL-62-Luc-TSHR cells. After 13 days, once tumors had engrafted and were palpable, 5 × 10⁶ healthy-donor T cells expanded for 7 days were injected intravenously either alone or together with M22-bsAb or K1-70-bsAb (0.5 mg/kg/dose). Mice then received bispecific antibody six times in total every 2–3 days and T cells from the same donor twice in total. Tumor volume was measured three times a week with a caliper and calculated as:

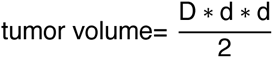

D is defined as the longer and d as the shorter side of the tumor.

### Flow cytometry

Flow cytometry was performed on a CytoFLEX™ (Beckman Coulter). Data were analyzed in FlowJo v10.0.1.

### Statistical analysis

If not stated otherwise, mean ± SD of technical replicate data is shown, number of replicates and donors are described in figure legends. Specific killing, LDH-release and IFN-γ-release differences between each construct and the corresponding control were assessed by ordinary two-way analysis of variance (ANOVA) with Dunnett’s multiple comparisons test. Thereby, anti-TSHR bsAbs were compared to allogeneic effects in target and effector co-culture (T+E) or anti-TSHR CAR T cells were compared to untransduced (UTD) T cells. Differences in the *in vivo* tumor growth model were determined by mixed-effects analysis (restricted maxium likelihood, REML) with Dunnett’s multiple comparisons test. GraphPad Prism v11 was used for statistical analysis and a two-sided p < 0.05 was considered significant.

## RESULTS

### TSHR expression is maintained in poorly differentiated thyroid cancer tissue and extracellular epitopes are accessible on fresh tissue

TSHR expression in DTC and ATC has been extensively studied, reporting stable expression in differentiated tumors and loss of expression upon anaplastic de-differentiation (7), but so far it remains understudied in PDTC (8). We therefore profiled TSHR expression by IHC on a TMA of 151 thyroid cancer cases (84 PTC, 29 FTC, 18 PDTC, 20 ATC) and 50 healthy thyroid samples, using the monoclonal anti-TSHR antibody EPR19751, which recognizes an extracellular epitope. PTC and FTC samples consistently showed strong membranous TSHR positivity. PDTC retained substantial TSHR expression at intensities comparable to PTC and FTC (Figure 1A, B). ATC samples were almost uniformly negative. We verified the accessibility of extracellular TSHR epitopes on native tumor tissue by flow cytometry using dissociated fresh tumor material and the K1-18 antibody (12) (Figure 1C). Together, these data establish that TSHR is preserved in patients with PDTC and that extracellular epitopes targetable by immunotherapy are accessible.

### Anti-CD3 X anti-TSHR bispecific antibodies derived from blocking or activating anti-TSHR antibodies can be produced at scale

To develop tumor-directed immunotherapy, we exploited available variable-region sequences from eight well-characterized anti-TSHR antibodies, four of human autoimmune origin (M22 (10, 40), K1-18 and K1-70 (12), 5C9 (11)) and four of murine origin (IRI-SAb2, IRI-SAb3 and 1H7 (41), 9D33 (29)), covering activating and blocking binding properties (Figure 2A). To generate CD3-engaging bsAbs, heavy- and light chain variable regions (VH/VL) were assembled to create tumor-binding Fab arms that were paired to the clinically validated anti-CD3 scFv via the human Fc region and produced in CHO-S cells (Figure 2B)(35). All eight antibody sequences yielded monomeric bsAbs in sufficient quantities for downstream functional characterization, with SDS-PAGE confirming the expected chain composition under reducing and non-reducing conditions (Supplementary Figure 1A–E). Using Jurkat-NFAT-Luc reporter cells co-cultured with TSHR-expressing target cells, we detected binding of five bsAbs (M22, K1-18, K1-70, 5C9 and 9D33) to both TSHR and CD3 (Figure 2C). The anti-TSHR antibody sequences from the bsAbs with confirmed TSHR and CD3 binding were then used as template to produce CARs with target binding scFvs in both VH–VL (HL) and VL–VH (LH) orientations using a second generation 4-1BB/CD3ζ signaling backbone. The Jurkat-NFAT-Luc reporter assay confirmed specific TSHR binding by the M22-HL, K1-18-HL, K1-70-HL and 5C9-LH CAR constructs, whereas the 9D33-LH CAR did not show a signal above control (Figure 2D).

**Figure 2.**
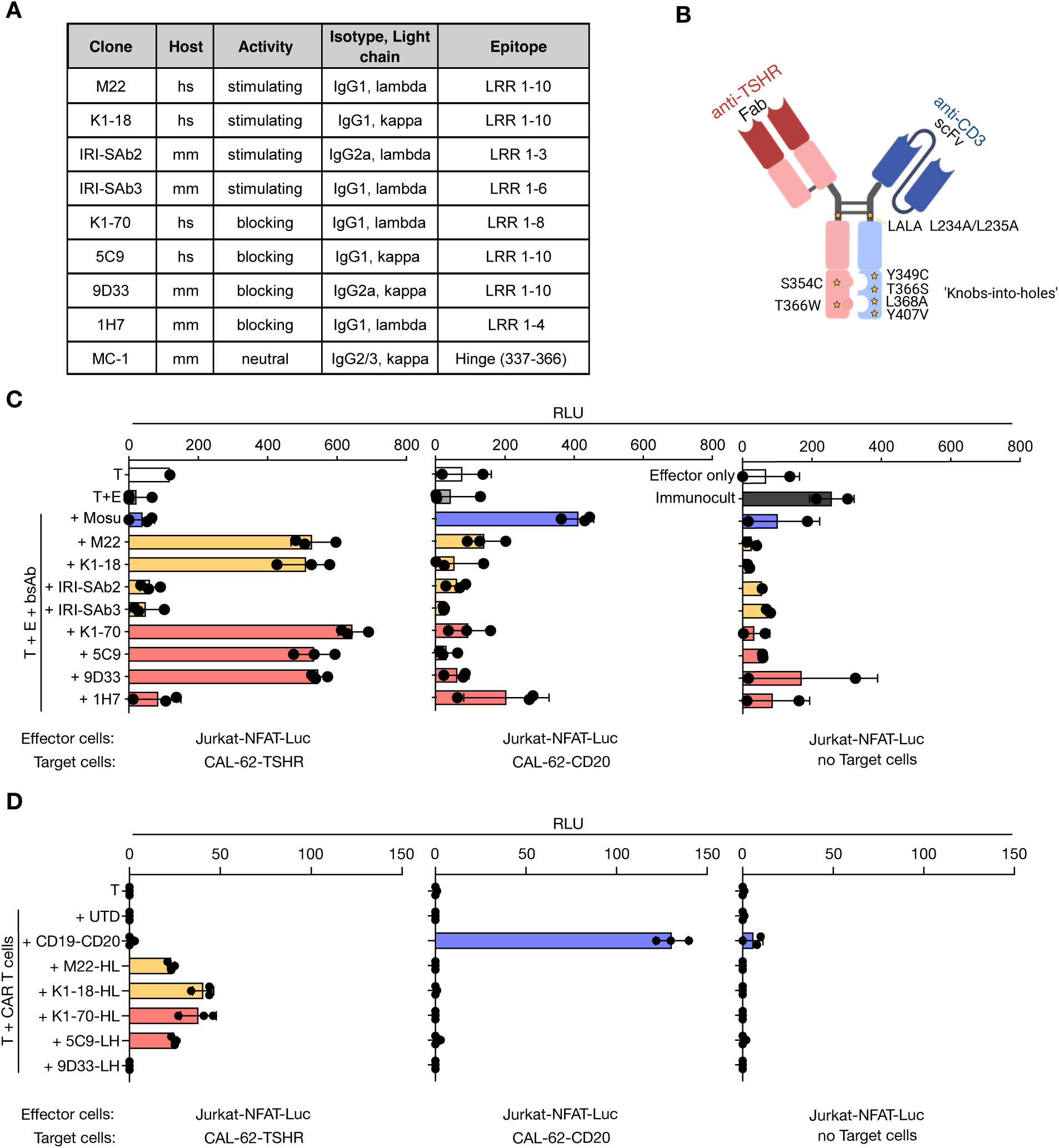
Generation and characterization of fully human anti-TSHR bispecific antibodies. **A** Panel of anti-TSHR antibodies used in this study, with original host, receptor-modulating activity (stimulating or blocking), isotype, light-chain usage and epitope. **B** Schematic depiction of the IgG-based CD3-engaging bispecific antibody (bsAb) format used for anti-TSHR bsAbs. The star symbols represent mutations introduced to the antibody. Created with BioRender.com. **C** Relative Light Units (RLU) of Jurkat-NFAT-Luc effector cells co-cultured for 24 h with indicated target cells in a 5:1 effector-to-target (E:T) ratio and 10 nM of anti-TSHR bsAb or mosunetuzumab (Mosu). Effector cells incubated with immunocult activation antibodies (1:50) served as control. **D** RLU of Jurkat-NFAT-Luc cells transduced with anti-TSHR CAR and incubated with indicated target cells (E:T = 5:1) for 24 h. CD19-CD20 CAR T cells and untransduced (UTD) T cells served as controls. Each dot represents one technical replicate with error bars as standard deviation (SD). Hs: Homo sapiens, mm: Mus musculus, LRR: leucine-rich repeat, RLU: relative light units, Mosu: mosunetuzumab, E: effector cells, T: target cells.

### Anti-TSHR bsAbs and CAR T cells selectively kill TSHR-expressing thyroid cancer cell lines *in vitro*

We next examined the cytolytic capacity of all anti-TSHR bsAbs and CAR T cells using GFP^+^ CAL-62-Luc-TSHR as target cells and healthy donor (HD) T cells as effector cells (Figure 3A–D). At an effector-to-target (E:T) ratio of 5:1 the bsAbs M22, K1-18, K1-70, 5C9 and 9D33 induced antigen-specific killing and reduced proliferation of TSHR-positive target cells accompanied by T cell activation and IFN-γ secretion (Figure 3B–D left). Among these, K1-70, M22 and K1-18 showed the strongest overall activity and were identified as the leading candidates (Figure 3B–D left). The same efficacies were observed using C643-Luc-TSHR cells as target cells (Supplementary Figure 2A, B). CAL-62-Luc-CD20 or C643-Luc-CD20 target cells lacking the TSHR did not exhibit any reduction in cell proliferation or viability and effector cells were not activated via bispecific antibody engagement above the allogeneic reaction in the co-culture (Figure 3 B–D, Supplementary Figure 2 A–C). Receptor modulation by the bsAbs was assessed in the CHO-TSHR-Luc reporter cell line. The stimulating clones M22 and K1-18 activated TSHR in the bsAb format to a degree comparable with their parental monospecific antibodies (msAb), and K1-70-bsAb blocked bovine TSH(bTSH)- or M22-msAb-induced stimulation in the same concentration range as K1-70-msAb indicating that reformatting preserved the receptor-modulating character of these clones (Supplementary Figure 2D– F). In contrast, 5C9- and 9D33-bsAb produced only limited blocking over the concentrations tested (Supplementary Figure 2D–F).

**Figure 3.**
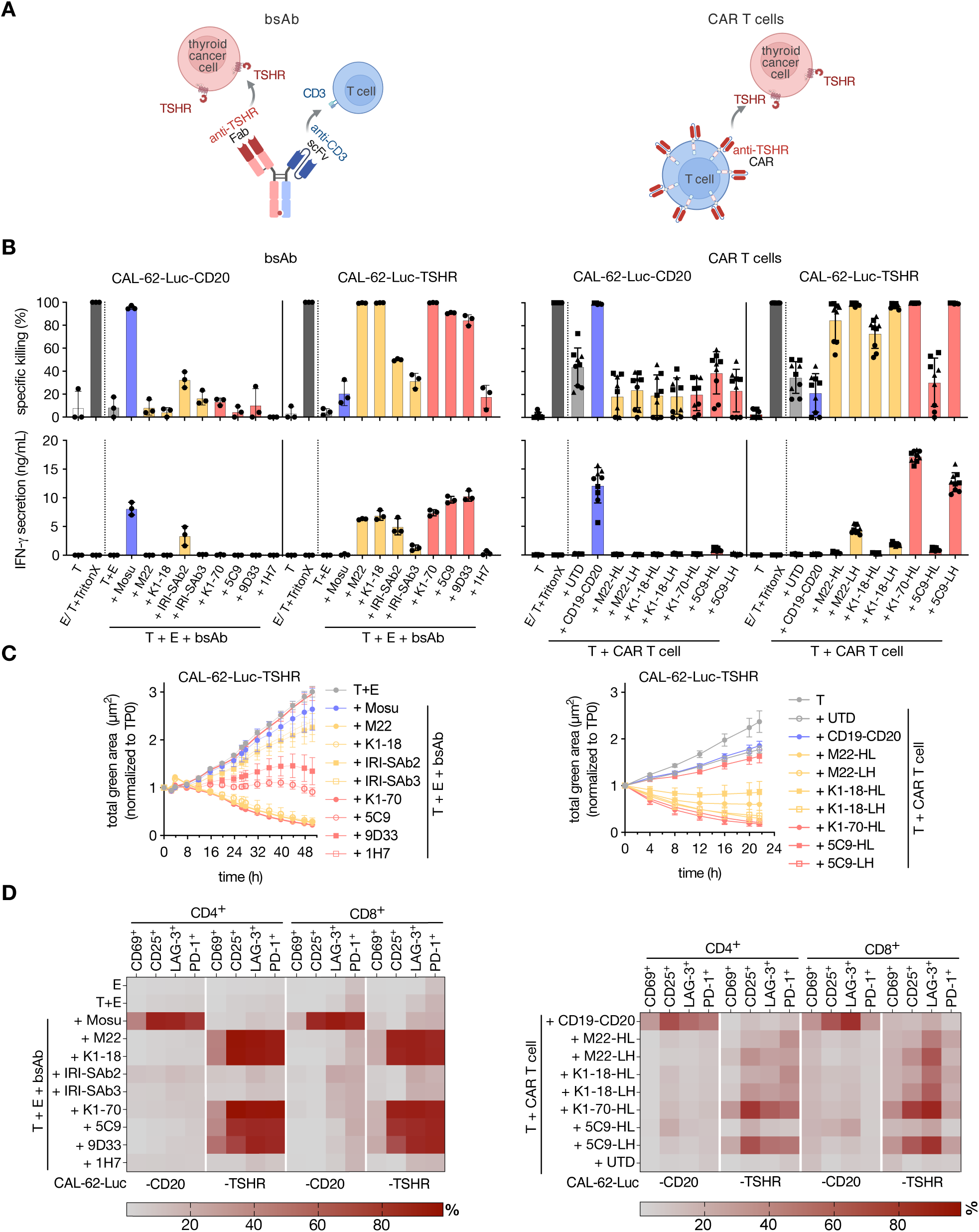
Selective killing of TSHR-expressing thyroid cancer cells by bsAbs and CAR T cells *in vitro*. **A** Schematic depiction of the two effector modalities. Left: the anti-TSHR bispecific antibody bridges TSHR on the tumor cell and CD3 on the T cell through an anti-TSHR Fab and an anti-CD3 scFv connected by a knob-into-hole Fc. Right: the same anti-TSHR scFv expressed on the T cell surface as a CAR, with hinge, transmembrane and 4-1BB/CD3ζ signaling domains. Created with BioRender.com. **B** Antigen-selective cytolysis of CAL-62-Luc-TSHR or CAL-62-Luc-CD20 cells by the eight anti-TSHR bsAbs at 10 nM (left) and by anti-TSHR CAR T cells in both scFv orientations (HL: VH–VL, LH: VL–VH), in co-culture with HD T cells at an E:T ratio of 5:1. Cytotoxicity was determined by luciferase read-out (top) and IFN-γ secretion (bottom) with three technical replicates for bsAbs and three biological replicates (●, ▴, ▪) with three technical replicates each for CAR T cell analyses. Error bars indicate the SD. Mosunetuzumab (Mosu) is the bsAb and UTD and CD19-CD20-CAR T cells are the CAR controls. **C** Proliferation of GFP⁺ target cells co-cultured as in B, monitored over 48 h (left) or 24 h (right) via live cell imaging normalized to time point = 0 (TP0). Each dot represents three technical replicates (left) or three biological and three technical replicates (n = 9, right) with error bars as SD. **D** Heat maps of activation markers (CD69, CD25, LAG-3, PD-1) on CD4⁺ and CD8⁺ T cells for bsAbs (left) and CAR T cells (right) with replicates described in B. E, effector cells; T, target cells.

Anti-TSHR CAR T cells incorporating clones from the validated bsAb subset showed comparable viability, transduction efficacy, CD4:CD8 ratios, and activation marker expression (Supplementary Figure 3A, B). At an E:T ratio of 5:1, they induced TSHR-specific killing, IFN-γ secretion and T cell activation, with K1-70-HL and 5C9-LH demonstrating the strongest activity (Figure 3B–D right). LDH-release assays corroborated the cytotoxicity rankings of both bsAbs and CAR T cells (Supplementary Figure 4A, B), while killing of TSHR-negative targets remained at the allogeneic baseline (Figure 3B–C, Supplementary Figure 3C). CAR activity depended on scFv chain orientation, supporting evaluations of both orientations when reformatting autoimmune antibodies into CARs (Figure 3B–D).

For the subsequent downstream analysis, candidates were narrowed down in both formats. Among the CAR constructs, K1-70-HL and 5C9-LH exhibited the strongest target cell killing and CAR T cell activation. For both constructs, the E:T ratio of 5:1 was optimal (Supplementary Figure 4D). Among the bsAbs, M22, K1-18 and K1-70 were the most potent. K1-70-bsAb was taken forward due to its blocking nature (Supplementary Figure 2E, F). To analyze a stimulating counterpart, we selected the M22-bsAb due to its high production yield (Supplementary Figure 1C). Titration of M22- and K1-70-bsAbs further demonstrated activity in the low-to sub-nanomolar range with killing saturation at 1 nM (Supplementary Figure 5A–G) and IC50 values 0f 0.07– 0.3 nM for K1-70 and 0.01–0.03 nM for M22 (Supplementary Figure 5E).

### Anti-TSHR bispecific antibodies control TSHR-positive tumors *in vivo*

To assess the therapeutic activity of bsAbs *in vivo*, NSG mice subcutaneously injected with CAL-62-Luc-TSHR xenografts were treated intravenously with human T cells together with M22-bsAb or K1-70-bsAb, or T cells alone over a period of two weeks (Figure 4A). Both bsAbs significantly suppressed tumor growth relative to T cells alone (Figure 4B) and suppression was consistent across individual animals in all three arms (Figure 4C). No overt toxicity, weight loss or behavioral abnormalities were observed in the treated mice.

**Figure 4.**
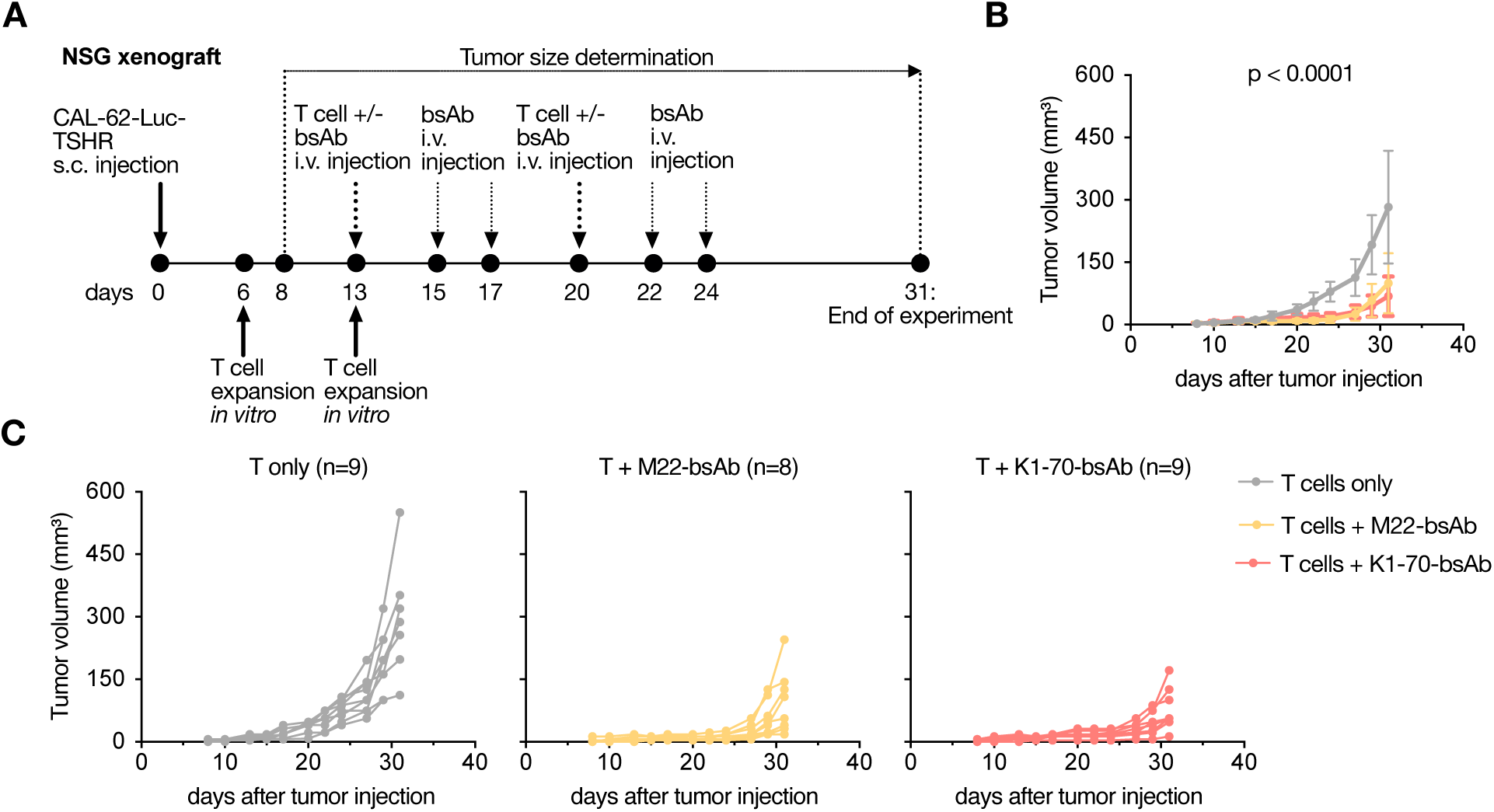
Effect of anti-TSHR bispecific antibodies in *in vivo* TSHR-positive xenografts. **A** Workflow of the NSG CAL-62-Luc-TSHR xenograft model. The T cells used for the first intravenous injection on day 13 had been activated and expanded for 7 days beforehand. The same 7-day activated and expanded T cell product from the same donor was used for both intravenous T cell injections. **B** Mean tumor volume over time for mice receiving T cells alone (n = 9), or with addition of M22-bsAb (n = 8), or K1-70-bsAb (n = 9). Each dot represents the mean of the respective biological replicates with error bars as SD. For statistical analysis, a mixed-effects model (REML) with Dunnett’s multiple comparisons test was used. **C** Individual tumor growth curves for each arm described in B.

### Patient-derived T cells mediate fully functional anti-TSHR bsAb and CAR T cell responses

Preserved cytolytic function of patient-derived T cells is crucial for successful clinical translation of autologous immunotherapies. To explore the feasibility of our approach in patient-derived T cells, peripheral blood T cells were isolated from two patients with PTC and one patient with FTC with a poorly differentiated component, all of whom had undergone surgery and/or radioiodine treatment. The cytolytic capacity of the patient-derived T cells was subsequently assessed in co-culture with CL-62-Luc-TSHR target cells in the presence of the two leading bsAb formats M22 and K1-70 and compared with that of T cells from our four HDs. Specific anti-TSHR bsAb-mediated killing, IFN-γ secretion and activation-marker up-regulation were of comparable magnitude in patient- and HD-derived T cells (Figure 5A, B; Supplementary Figure 6A, B). In addition, we produced 5C9-LH- or K1-70-HL-CAR T cells using T cells originating from three HDs and three patients with PTC, FTC, or PDTC after surgery and/or radioiodine treatment. HD- and patient-derived CAR T cells exhibited comparable viability and transduction efficiencies over three weeks of expansion (Supplementary Figure 7A), together with almost identical activation marker expression (Supplementary Figure 7B). Although patient-derived CAR T cells exhibited a shift towards higher CD4:CD8 ratios (Supplementary Figure 7A), their antigen-specific cytotoxicity against CAL-62 Luc TSHR target cells was comparable to that of HD-derived CAR T cells, despite donor-to-donor variability (Figure 5C, D; Supplementary Figure 8A, B).

**Figure 5.**
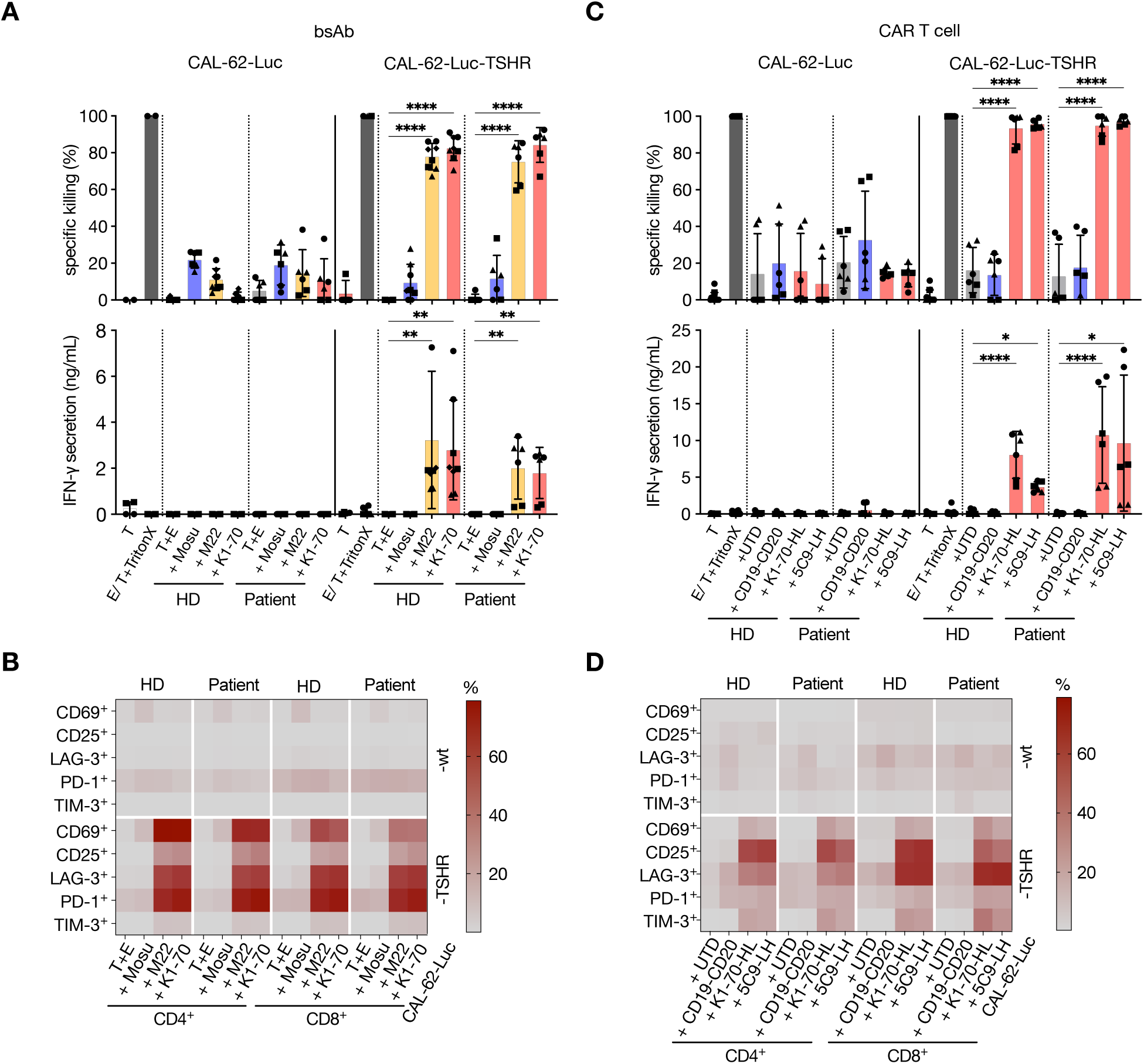
Functionality of bsAb and CAR T cell treatment with patient-derived T cells. **A** Cytolysis of CAL-62-Luc-TSHR versus parental CAL-62-Luc cells by 1 nM anti-TSHR M22- and K1-70-bsAb in co-culture with T cells from HD or from thyroid cancer patients. Cytotoxicity was determined by luciferase assay (top) and IFN-γ secretion (bottom) with mosunetuzumab (Mosu) serving as control. Per biological replicate indicated by different symbols (●, ▴, ▪, ◆), two technical replicates were used with error bars as SD. **B** Expression of activation markers (CD69, CD25, LAG-3, PD-1, TIM-3) on CD4⁺ and CD8⁺ T cells for the co-culture described in A. **C** Cytolysis of the same targets by anti-TSHR K1-70-HL- and 5C9-LH-CAR T cells generated from HD or patient T cells, with UTD and CD19-CD20 CAR T cells as controls. Per biological replicate indicated by different symbols (●, ▴, ▪), two technical replicates were used with error bars as SD. **D** Expression of activation markers for the co-culture in C. wt: parental CAL-62-Luc, E: effector cells, T: target cells. ****: p < 0.0001, **: p ≤ 0.01, *: p ≤ 0.05.

### Anti-TSHR bsAbs and CAR T cells eliminate target cells with low TSHR expression and across heterogeneous antigen distributions, with measurable bystander activity

TSHR expression levels on PDTC are largely underexplored. To investigate whether reduced surface TSHR expression or a lower proportion of TSHR-positive cells impairs bsAb- and CAR T cell-mediated killing, we engineered corresponding CAL-62-Luc target cell models. First, we generated two cell lines with stably reduced surface TSHR expression, CAL-62-Luc-IRESv7-TSHR and IRESv3-TSHR, by mutating the 11^th^ and 12^th^ AUG codons of the IRES element, respectively, with non-AUG triplets (Figure 6A, Supplementary Figure 9A). In another model, both lower levels of overall TSHR expression and variable frequencies of cells expressing TSHR was established making use of a doxycycline-inducible Tet-On TSHR construct (Figure 6A, Supplementary Figure 9B).

**Figure 6.**
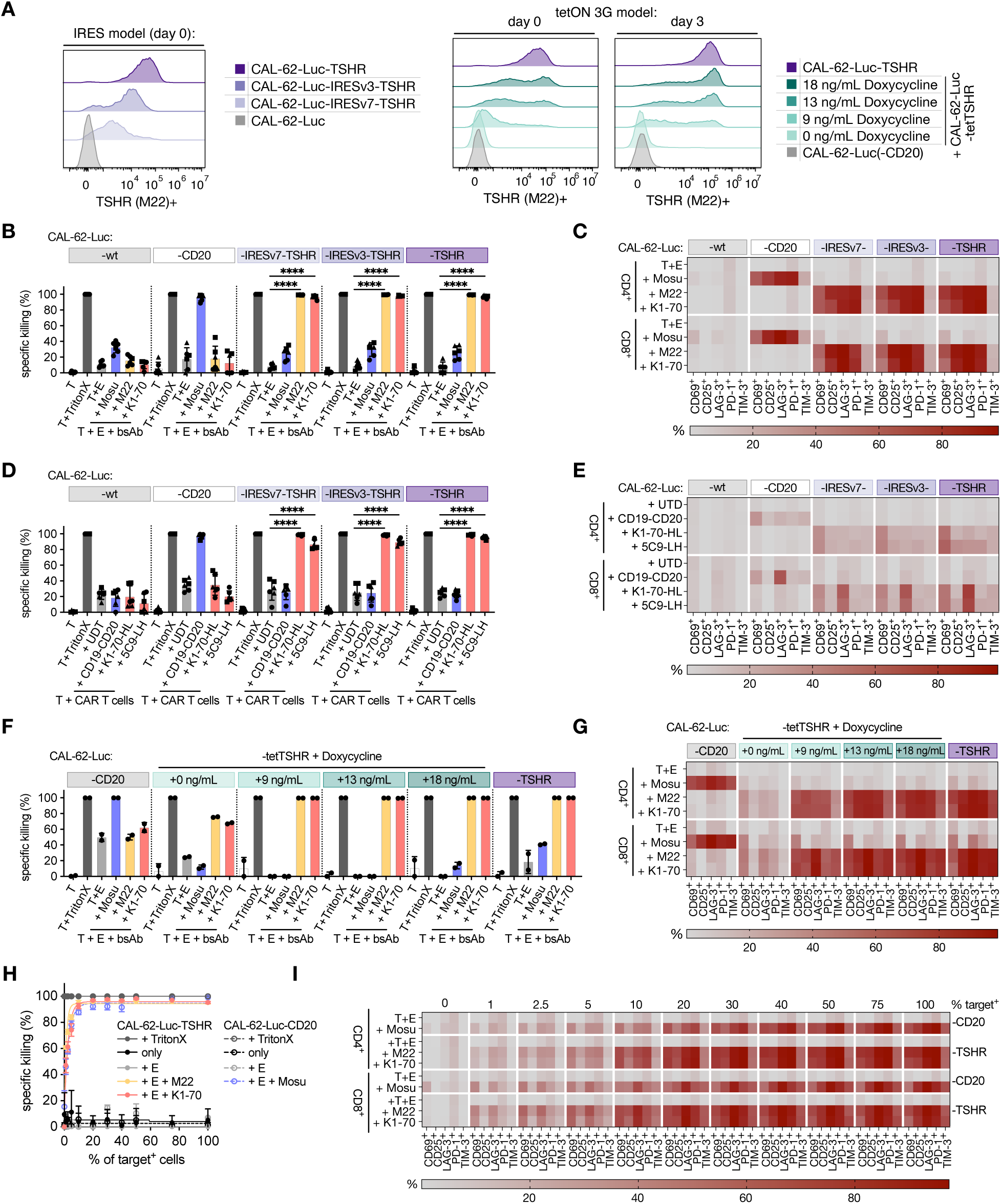
Influence of anti-TSHR bsAbs and CAR T cells on cells with reduced or heterogeneous antigen expression. **A** Flow-cytometric detection of surface TSHR (M22) on CAL-62-Luc/ -TSHR/ -IRESv3-TSHR/ -IRESv7-TSHR/ -CD20 or CAL-62-Luc-tetTSHR treated with 0 – 18 ng/mL doxycycline on day 0 or day 3 of co-culture and normalized to CAL-62-Luc-TSHR. **B** Specific killing of CAL-62-Luc-wt/ -CD20/ -IRESv7-TSHR/ - IRESv3-TSHR/ -TSHR target cells after co-culture with 1 nM of anti-TSHR M22- and K1-70-bsAb and HD T cells with mosunetuzumab (Mosu) as a control. Three biological replicates (●, ▴, ▪) were used with two technical replicates each. For statistical analysis, ordinary two-way ANOVA with Dunnett’s multiple comparisons test was used with error bars as SD. **C** Expression of activation markers on CD4⁺ and CD8⁺ T cells for the co-culture described in B. **D** Specific killing of the target cells in (A) by anti-TSHR K1-70-HL- and 5C9-LH-CAR T cells, with untransduced (UTD) and CD19-CD20-CAR T cells as controls. A total of three biological replicates as indicated by the symbols (●, ▴, ▪) were used with two technical replicates each. Ordinary two-way ANOVA with Dunnett’s multiple comparisons test was used for statistical analysis with error bars as SD. **E** Corresponding heat map of activation markers of the co-culture performed in D. **F** Specific killing of CAL-62-Luc-tetTSHR cells induced with 0, 9, 13 or 18 ng/mL doxycycline, with CAL-62-Luc-CD20 and constitutively TSHR-expressing CAL-62-Luc-TSHR cells as references. Each dot represents one technical replicate with error bars as SD. **G** Corresponding expression of activation markers. **H** Specific killing of mixed co-cultures containing defined fractions (0–100%) of antigen-positive cells CAL-62-Luc-TSHR/ -CD20 mixed with antigen-negative CAL-62-Luc and treated with 1 nM of M22-bsAb, K1-70-bsAb or mosunetuzumab (Mosu) and HD T cells. Each dot represents the mean of three technical replicates with error bars as SD. **I** Expression of activation markers across the mixture series described in H. E, effector cells; T, target cells. ****: p < 0.0001.

We stained all cell line models by immunohistochemistry and compared their antigen expression levels with those in the patient TMA described in Figure1, placing the antigen densities tested *in vitro* in the context of patient tissue (Supplementary Figure 9C).

In the first model, M22- and K1-70-bsAbs as well as K1-70- and 5C9-CAR T cells eliminated CAL-62-Luc-IRESv3-TSHR and CAL-62-Luc-IRESv7-TSHR cells with efficacies comparable to those against control cells with high TSHR expression (Figure 6B–E). Effector cells also showed similar activation marker profiles following exposure to targets with reduced or high TSHR expression (Figure 6C, E).

For investigating the cytotoxicity in the TSHR-expression-inducible model, CAL-62 Luc tetTSHR cells were exposed to 0–18 ng/mL doxycycline for 72 h generating target populations with varying surface TSHR expression levels and proportions of TSHR-positive cells (Figure 6A, Supplementary Figure 9B). In 48 h co-cultures with these targets, bsAb-mediated cytotoxicity was preserved across the full doxycycline titration, with appreciable killing even at the lowest induction levels tested (Figure 6F). Activation marker profiles followed a similar pattern across induction levels (Figure 6G).

We next modeled intratumoral antigen heterogeneity at the population level using defined mixtures of antigen-positive CAL-62-Luc-TSHR or CAL-62-Luc-CD20 cells and antigen-negative CAL-62-Luc cells. Mixtures containing increasing proportions of antigen-positive targets were incubated with HD T cells and M22 or K1-70 anti-TSHR bsAbs, or with mosunetuzumab for the CD20 model (Figure 6H). Anti-TSHR bsAbs induced killing that substantially exceeded the proportion of TSHR-positive cells. Target cell death reached about 50% when only 1% of target cells carried TSHR and approximately 80% at 5% TSHR positivity and plateaued above 90% when approximately 10% or more of target cells expressed TSHR (Figure 6H). Mosunetuzumab showed a near-identical relationship between antigen-positive cell fraction and killing in mixed CD20-positive and CD20-negative cultures (Figure 6H). Surface expression of activation markers on effector cells increased with the proportion of antigen-positive targets (Figure 6H). Cytotoxicity patterns in all cell models were additionally confirmed by LDH release assay (Supplementary Figure 10, A–H).

Taken together, our data suggest that cytolysis is preserved at substantially reduced antigen densities and heterogenous TSHR-antigen expression within the tumor. Killing beyond the antigen-positive cell fraction is consistent with an amplifying bystander effect that is also a general property of clinically used CD3-engaging bsAbs.

## DISCUSSION

Patients with PDTC carry one of the highest unmet medical needs in thyroid oncology (3, 42). The natural history of PDTC is heterogeneous, but once RAI fails and TKI with or without immune checkpoint inhibition is exhausted, options reduce to chemotherapy of marginal benefit and to rare, genotype-defined treatments (BRAF, RET, NTRK) (4–6). Because PDTC retains substantial molecular continuity with DTC, including the TSHR-axis transcriptional program, it is in principle accessible to antigen-directed therapy, yet the field has lacked a tumor-selective immune-based strategy that does not depend on mutation-defined neoantigens. We took advantage of two convergent resources. TSHR is a near-ideal tumor-associated antigen for thyroid cancer with tightly restricted physiological expression and pharmacologically replaceable function, and the human anti-TSHR autoantibody repertoire offers sequence-defined binders with two decades of characterization in autoimmune thyroid disease (13–15). We confirmed TSHR expression in a clinically meaningful subset of PDTCs, generated eight anti-TSHR bsAbs, and used clones showing TSHR binding and cytolytic activity to develop CAR T cells in both HL and LH scFv orientations. While both treatment formats achieve potent, antigen-selective cytolysis *in vitro*, the leading bsAbs could further control TSHR-positive xenografts *in vivo*. Interestingly, the scFv chain orientation turned out to be decisive for CAR T cell function. T cells derived from patients with DTC or PDTC obtained during or after treatment retained full effector competence, providing first evidence of their suitability for both bsAb and CAR T cell approaches. Finally, we show that TSHR-directed immunotherapy retains cytotoxic activity against thyroid cancer cells with reduced or heterogenous antigen expression. The anti-TSHR antibody repertoire presented here provides fully human, sequence-defined variable regions for off-the-shelf bsAb candidates that retain parental receptor-modulating activity.

Accordingly, the central advance here is an off-the-shelf approach to TSHR targeting alone. To our knowledge these are the first peer-reviewed data for a CD3-engaging bispecific antibody against TSHR. Previous TSHR-directed efforts focused exclusively on DTC and CAR T cell approaches requiring patient-specific manufacturing (22, 23, 25). These studies also evaluated only a few antibody candidates, limiting comparative assessment (22–24). As off-the-shelf agents, bsAbs avoid patient specific manufacturing delays, allowing prompt treatment initiation and flexible dosing that can be adjusted, paused, and re-escalated (43).

These results refine the picture from prior work. The TSHR field has long debated whether TSHR remains expressed in dedifferentiated thyroid cancer at therapeutically meaningful levels (7, 8). Our IHC and flow cytometry data consolidate the view that PDTC, but not ATC, retains surface-accessible TSHR. Further, efforts to improve TSHR expression in patients are explored (44).The work that described the K1-18, K1-70, M22, 5C9, IRI-SAb2, IRI-SAb3 and 1H7 antibodies provided the sequences we now repurpose for tumor targeting (10–12, 40, 41). Across all clones we tested, five bsAbs mediated antigen-selective cytolysis across both stimulating and blocking parental clones, suggesting that target recognition is more relevant to effector-mediated killing than the parental signaling profile. Reporter-cell assays showed that reformatted bsAbs could stimulate TSHR or inhibit bTSH- and antibody-induced signaling, but preservation of their full parental signaling profiles was not comprehensively assessed. Stimulating bsAbs could therefore activate TSHR before engaging effector cells, allowing receptor signaling without cytolysis. In CAR T cells, target recognition is directly coupled to the effector cell itself.

The strong dependence of CAR T cell function on scFv chain orientation observed across multiple clones, argues that both VH–VL and VL–VH arrangements should be routinely screened when repurposing autoimmune-derived antibody sequences into a CAR backbone. The Fab-based bsAb format avoids this orientation choice, simplifying one aspect of construct design. K1-70-derived constructs perform best in both treatment formats, consistent with the broad ectodomain engagement of K1-70, identifying this clone as the lead candidate for clinical development (13). Cytolytic activity is retained at substantially reduced surface TSHR levels demonstrated in mutated IRES and Tet-induced cell line models. Retained cytotoxicity at reduced TSHR expression supports further evaluation in PDTC, in which antigen density is expected to be both lower and more heterogeneous than in DTC, and is consistent with the antigen-density thresholds described for CAR constructs (7, 8, 45). However, our engineered models do not establish a clinical expression threshold and short-term killing at low antigen density does not necessarily predict sustained tumor control *in vivo*. Sensitivity to low antigen levels may also increase activity against normal TSHR-expressing tissues. Recent TSHR-directed CAR experiments reported transient thyroid-follicle injury in mice reinforcing the need to assess normal-tissue safety (22).

The preserved cytolytic function in patient-derived T cells supports the feasibility of both formats, aligning with what has been described for CAR T cell manufacturing from heavily pretreated hematological patients, and thus reducing a key translational risk (43, 46).

An observation that emerged consistently from our mixed-population experiments is that anti-TSHR bsAb-mediated cytolysis is not strictly antigen-restricted. In co-cultures with defined fractions of TSHR-positive and TSHR-negative cells, both M22- and K1-70-bsAb killed far more cells than carried the antigen. Roughly half of the culture was lysed when only 1% of cells were TSHR-positive, and killing plateaued above 90% from about 10% positive cells upwards. This bystander activity was not unique to anti-TSHR constructs as the clinically approved anti-CD20 x anti-CD3 bsAb produced a bystander effect of comparable magnitude in mixed CD20-positive/ CD20-negative co-cultures. This is consistent with the effect being a property of the CD3-engager modality rather than of any specific target antigen, as previously reported for BiTE constructs (47). Even though we did not investigate this further, possible candidate mechanisms include soluble effector mediators, such as IFN-γ, tumor necrosis factor (TNF) or perforin and granzymes, and damage-associated factors released from lysing antigen-positive cells (47, 48). Fas–FasL-dependent killing of antigen-negative cells has also been shown to contribute substantially in T cell immunotherapy (47, 49). Taken together, these data establish the anti-TSHR bispecific antibody as an off-the-shelf treatment candidate for TSHR-positive thyroid cancer, with the autologous CAR format as the alternative where it is warranted. Both, built around the K1-70 binder, achieve potent, antigen-selective elimination of TSHR-positive thyroid cancer cells under conditions relevant to PDTC. Anti-TSHR bsAbs merit early clinical evaluation as an off-the-shelf modality in progressive DTC and PDTC after treatment failure. Their manufacturability, the absence of T cell engineering, and compatibility with concurrent systemic therapy furthermore highlight their potential. Beyond thyroid cancer, this work demonstrates that organ-specific autoantibody repertoires can be systematically repurposed as a sequence-defined discovery resource for tissue-restricted tumor antigens.

## Acknowledgements

The authors thank the patients and healthy donors who contributed samples to this work, and the staff of the participating clinical departments for assistance with sample collection. We thank Adeline Stiefvater and Anja Muskulus for excellent technical support and Mihaela Barbu-Stevanovic, Cécile Cumin, Marion Wernli, Andrea Mazzoleni and Morgane Hilpert from the flow cytometry core facility at the Department of Biomedicine at the University Hospital Basel for excellent technical assistance.

## Authors’ contributions

Conception and design of the study: M.B., C.F., H.L., M.P. Provision of clinical samples and patient data: K.L., J.J., L.P.M., R.K.-W., B.K., C.R. Pathology review and IHC analysis: M.Ba., A.W., C.W. Provision of critical reagents and constructs: M.P., R.L., T.F.D., E.D’A. Methods development and experimental work: C.F., A.E., B.B, M.Bu., F.F., C.S. Data analysis and interpretation: C.F., A.E., H.L., M.B. Drafting of the manuscript: C.F., M.B., A.E. Critical revision of the manuscript for important intellectual content: all authors. Supervision and funding acquisition: M.B. All authors approved the final version of the manuscript and agree to be accountable for all aspects of the work.

During the preparation of this manuscript, the authors used Claude Opus 5.5 to improve the language and readability of the text. The tool was not used to generate scientific content, analyze data, or interpret results. The authors reviewed and edited all output and take full responsibility for the content of the publication.

## Availability of data and material

All data supporting the findings of this study are presented in the manuscript and its supplementary materials. Raw flow-cytometry files, IHC scans, *in vivo* datasets and source data for all plots are available from the corresponding author on reasonable request. The variable-region sequences of the anti-TSHR antibodies used in this study are available in the cited primary literature.

## List of abbreviations

ANOVA: analysis of variance;
ATC: anaplastic thyroid carcinoma;
BL: baseline;
bsAb: bispecific antibody;
bTSH: bovine thyroid-stimulating hormone;
CAR: chimeric antigen receptor;
CHO: Chinese hamster ovary;
DAPI: 4′,6-diamidino-2-phenylindole;
DTC: differentiated thyroid carcinoma;
E:T: effector-to-target ratio;
FFPE: formalin-fixed, paraffin-embedded;
FTC: follicular thyroid carcinoma;
GFP: green fluorescent protein;
HD: healthy donor;
IFN-γ: interferon gamma;
IHC: immunohistochemistry;
IRES: internal ribosome entry site;
LDH: lactate dehydrogenase;
MFI: median fluorescence intensity;
msAb: monospecific antibody;
NFAT: nuclear factor of activated T cells;
NSG: NOD/SCID/IL2Rγnull mouse;
PDTC: poorly differentiated thyroid carcinoma;
PTC: papillary thyroid carcinoma;
qPCR: quantitative polymerase chain reaction;
RAI: radioactive iodine;
REML: restricted maximum likelihood;
RLU: relative light unit;
scFv: single-chain variable fragment;
SD: standard deviation;
SDS-PAGE: sodium dodecyl sulfate–polyacrylamide gel electrophoresis;
SFFV: spleen focus-forming virus;
TKI: tyrosine kinase inhibitor;
TMA: tissue microarray;
TNF: tumor necrosis factor;
TP0: time point 0;
TSHR: thyroid-stimulating hormone receptor;
UTD: untransduced;
wt: wild-type.

