## Supplementary_Material for "Targeting poorly differentiated thyroid cancer with bispecific antibodies and CAR T cells from the human anti-TSHR autoantibody repertoire"

### **Supplementary Methods**

#### **Immunohistochemistry**

Staining was performed with the ZytoChem Plus (AP) Polymer kit (Leica, DS9800) on a fully automated Leica Bond stainer according to the manufacturer's instructions.

#### **Dissociation of fresh tumor tissue**

Tissue was digested in 1× Accutase (Innovative Cell Technologies) diluted 1:2 in high-glucose DMEM (Thermo Fisher Scientific), supplemented with 1 mg/mL collagenase IV (Worthington), 1 mg/mL hyaluronidase (Sigma), 0.01 KU/mL DNase I type IV (Sigma) and 10 μM Y-27632 (Stem Cell Technologies) at 37 °C with shaking at 120 rpm.

#### **Bispecific antibody constructs**

The anti-CD3 arm consists of the UCHT1 variable heavy and light chains separated by a 15-amino-acid linker, followed by a 4GS linker and the IgG1 heavy-chain constant region with a truncated hinge (EPKSCDKTHTCPPCP). All TSHR- and CD3-targeting expression constructs were cloned into pcDNA3(+) under the control of the CMV promoter and preceded by the IL-2 leader sequence.

#### **Bispecific antibody production and purification**

FreeStyle™ CHO-S cells were transfected using the MaxCyte STx™ electroporation system and cultured at 37 °C, 5% CO<sub>2</sub>, 120 rpm in CD Opti-CHO medium supplemented with 1% HT supplement, 1% GlutaMAX, 1% Pluronic F-68 and 15% CHO CD EfficientFeed™ A. Sodium butyrate was added to 1 mM 24 h after electroporation, the temperature was shifted to 32 °C, and 10% culture volume of feed stock (69.5% CHO CD EfficientFeed™ A, 13.9% Bacto™ TC Yeastolate, 3.5% GlutaMAX, 6.2% glucose, 6.9% Gibco™ Function MAX™ Titer Enhancer) was added every three days. After at most 14 days, or once viability fell below 50%, supernatant was harvested, centrifuged for 15 min at 2,000 rpm and 4 °C, filtered through a 0.2 μm Filtropur and a Sartoclear Dynamics® Lab 0.22 μm depth filter, and stored at –20 °C. Supernatants diluted 1:1 in Tris pH 8.0 were loaded for affinity capture; chromatography was run on an ÄKTA pure™ 150 T system (Cytiva).

#### **Lentiviral production**

Virus particles were produced in Lenti-X 293T cells (Takara) transfected by CaCl<sub>2</sub> with the third-generation packaging plasmids pMDLg/pRRE, pRSV-Rev and pVSV-G together with the transfer vector. Supernatants were concentrated by PEG-8000 precipitation, resuspended in DMEM medium supplemented with GlutaMAX™, 10% heat-inactivated FBS, 100 U/mL penicillin and 100 μg/mL streptomycin) used for target cells or in T cell medium (RPMI-1640 with GlutaMAX™, 10% heat-inactivated human serum, 100 U/mL penicillin, 100 μg/mL streptomycin, 50 μM 2-mercaptoethanol) supplemented with 50 IU/mL IL-2 (StemCell Technologies) and 10 ng/mL IL-15 and IL-7 (PeproTech) for CAR T cells. Virus was stored at –80 °C and titrated by qPCR (qPCR Lentivirus Titer Kit, abm).

#### **CAR T cell generation and flow-cytometry panel**

Transduction was supported by 0.5 mg/mL Synperonic F108 (Sigma-Aldrich). Cells were expanded in PRIME XV T-Cell CDM medium (FUJIFILM Irvine Scientific) with 10 ng/mL IL-15 and IL-7 in G-Rex 24-well plates (Wilson Wolf). The anti-G4S-linker (Cell Signaling Technologies, Cat# 71645S) with secondary anti-rabbit IgG AlexaFluor488 (Cell Signaling Technologies, Cat# 4412S) was used to measure transduction efficacy. With APC anti-human CD69 (FN50, RRID:AB\_314845, Cat# 310910), BV510 anti-human CD25 (M-A251, RRID:AB\_2563874, Cat# 356120), PE/Cyanine7 anti-human CD223 (LAG-3, 11C3C65, RRID:AB\_2629753, Cat# 369310), PE anti-human CD279 (PD-1, EH12.2H7, RRID:AB\_940483, Cat# 329906), APC/Cyanine7 anti-human CD366 (TIM-3, F38-2E2, RRID:AB\_2565717, Cat# 345026), BV605 anti-human CD4 (SK3, RRID:AB\_2734348, Cat# 344646) and BV650 anti-human CD8 (SK1, RRID:AB\_2564510, Cat# 344730) (all BioLegend) antibodies, T cell activation was determined. For viability, the DAPI staining solution (Mitenyi) was used.

#### Calculation of specific killing and cytotoxicity

$$\% \text{ specific killing} = 100 \times ((RLU_{\text{sample}} - RLU_{\text{target-only}}) / (RLU_{\text{TritonX}} - RLU_{\text{target-only}}))$$

$$\% \text{ Cytotoxicity (LDH release)} = 100 \times (((Absorbance_{\text{sample}} - Absorbance_{\text{effector-only}}) - Absorbance_{\text{target-only}}) / (Absorbance_{\text{TritonX}} - Absorbance_{\text{target-only}}))$$

The effector-only term was set to 0 when no effector cells were added. Negative values were set to 0.

#### ***In vivo* xenograft model following the ARRIVE 2.0 guidelines**

Animal and husbandry: Twenty-six male NSG mice from in-house breeding at the animal core facility of the University Basel were used. Mice were 7-9 weeks of age with a mean body weight of 28 g at the start of treatment. Animals were maintained under specific-pathogen-free conditions, housed in individually ventilated cages at a maximum of five mice per cage, at room temperatures on a 12:12 hour light:dark cycle and with nesting material and shelter as environmental enrichment. Only male mice were used, based on availability from in-house breeding.

Study design and allocation: Mice were injected subcutaneously (s.c.) into the right flank under isoflurane inhalation anesthesia. On the day of first treatment, tumor volume was measured and mice were ranked by baseline tumor volume. Animals were allocated to three treatment arms by size matching to ensure comparable baseline tumor burden across arms leading to the three treatment groups T cells only (n = 9), T cells and M22-bsAb (n = 8) and T cells and K1-70-bsAb (n = 9). The individual mouse was the experimental unit. Group sizes were based on prior experience with this xenograft model and kept to the minimum, in line with the 3Rs.

Procedures: T cells from all arms derived from the same donor. No post-procedural analgesia was administered, as s.c. was not expected to cause lasting discomfort. T cells and bsAbs were administered intravenously (i.v.). after mice were placed under red light for 5 – 15 minutes. Mice were shortly placed in a restrainer, injected in the tail vein and observed in their cages for 5 – 10 minutes post injection.

Outcome measures and monitoring: Tumor volume was determined with a caliper. No blinding could be performed. Predefined humane endpoints were clinical symptoms including tumor ulceration or tumor volume exceeding 1500 mm<sup>3</sup>. No animal reached the humane endpoint before day 31. Mice were euthanized by CO<sub>2</sub> inhalation.

Inclusion and exclusion criteria: All 26 animals established palpable tumors, were allocated to treatment, completed the study to day 31 and were included in the analysis. No animals were excluded and no data points were removed.

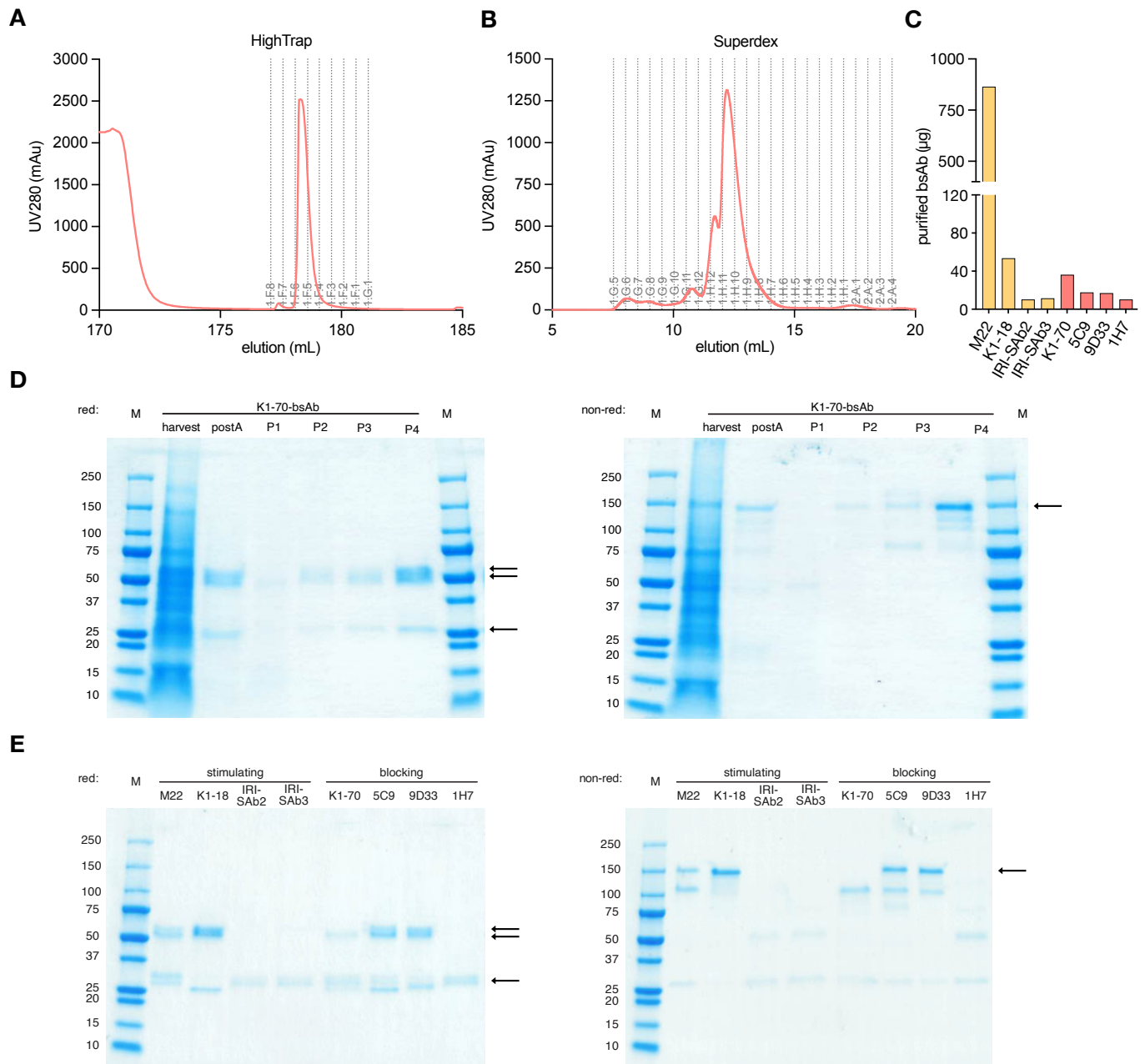

**Supplementary Figure 1. Production and purification of anti-TSHR bispecific antibodies.** **A** Exemplary elution and fractionation profile obtained by purification of FreeStyle™ CHO-S cell supernatant on a HiTrap™ Fibro PrismA column. **B** Exemplary elution and fractionation profile obtained by separating protein A-purified anti-TSHR bsAb on a Superdex 200 Increase 10/300 column. **C** Amount of purified, monomeric bsAb obtained after a typical production starting from 80 million FreeStyle™ CHO-S cells transfected with 120 µg bsAb-encoding plasmid DNA. Column colors indicate whether the bsAb derives from a stimulating (yellow) or blocking (red) anti-TSHR antibody. **D** SDS-PAGE analysis of purification fractions of K1-70-bsAb (131 kDa) under reducing (left) and non-reducing (right) conditions. For the former, the bsAb is separated into its anti-TSHR light chain (25.5 kDa), anti-TSHR heavy chain (51.5 kDa) and anti-CD3-scFv-Fc chain (54 kDa). Samples were taken after supernatant harvesting, HiTrap™ Fibro PrismA (postA) and different Superdex peaks (P1-P4). **E** SDS-PAGE analysis of anti-TSHR bsAbs under reducing (left) and non-reducing (right) conditions.

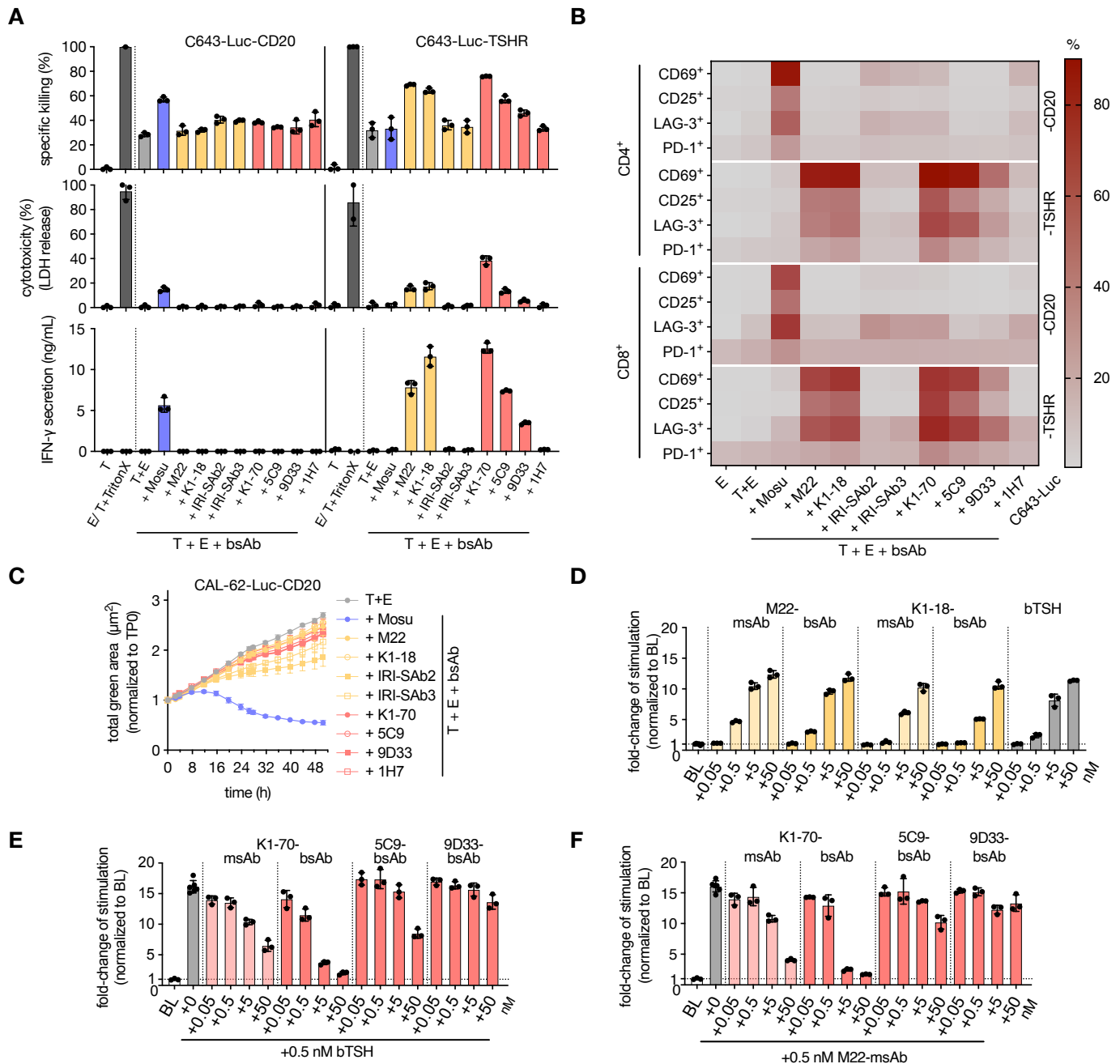

**Supplementary Figure 2. Anti-TSHR bispecific antibodies selectively kill TSHR-expressing thyroid cancer cells in a second target cell line, and retain the receptor-modulating properties of their parental clones. A** Antigen-dependent cytotoxicity of C643-Luc-TSHR (target, right) or C643-Luc-CD20 (control, left) cells by the eight anti-TSHR bsAbs (10 nM) after 24 h co-culture with HD CD3<sup>+</sup> T cells at an effector-to-target (E:T) ratio of 5:1. Specific killing (%) by luciferase assay (top), cytotoxicity by lactate dehydrogenase (LDH) release (middle) and IFN-γ secretion (ng/mL) (bottom) are shown. Each dot represents one technical replicate with error bars as SD. Color coding as in Supplementary Figure 1C. **B** Heat map of activation markers (CD69, CD25, LAG-3, PD-1) on CD4<sup>+</sup> and CD8<sup>+</sup> T cells in the same co-culture. Each square represents the mean of three technical replicates. **C** Proliferation of GFP<sup>+</sup> CAL-62-Luc-CD20 control target cells during co-culture with the bsAbs and HD CD3<sup>+</sup> T cells, monitored by live-cell imaging (Incucyte S3) over 48 h and expressed as total green area normalized to time point = 0 (TP0). Each dot represents the mean of three technical replicates with error bars as SD. **D** TSHR stimulation in CHO-TSHR-Luc reporter cells exposed to the parental monospecific antibodies (msAb) M22 and K1-18 or the corresponding bsAbs at 0.05–50 nM, normalized to baseline (BL). BL is defined as the luciferase activity of CHO-TSHR-Luc cells without any stimulators. Bovine TSH (bTSH) is the reference agonist. **E** Blocking of bTSH-induced TSHR stimulation in CHO-TSHR-Luc reporter cells by K1-70-msAb, K1-70-bsAb, 5C9-bsAb and 9D33-bsAb over 0.05–50 nM, normalized to BL. **F** Blocking of M22-msAb-induced TSHR stimulation by

antibodies mentioned in E. In D, E and F, each dot represents one technical replicate with error bars as SD. T: target cells, E: effector cells

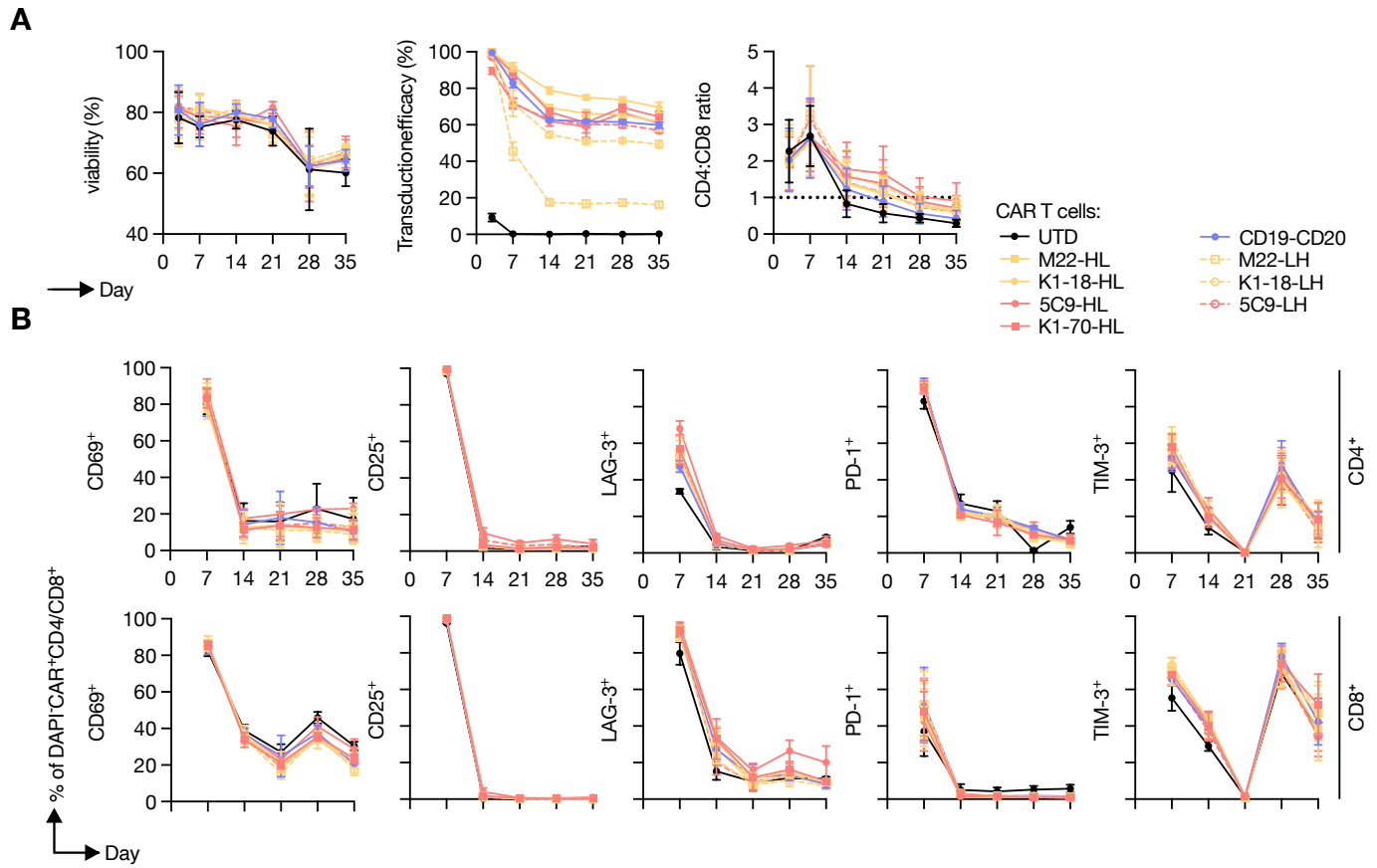

**Supplementary Figure 3. Characterization of the anti-TSHR CAR T cells.** **A** Viability (left), transduction efficiency (middle) and CD4:CD8 ratio (right) of HD CAR T cells observed over a period of 35 days. Constructs are the anti-TSHR CARs in both scFv orientations (HL: VH–VL; LH: VL–VH) together with CD19-CD20-CAR and untransduced (UTD) control cells. **B** Activation markers (CD69, CD25, LAG-3, PD-1 and TIM-3) on DAPI<sup>-</sup>CAR<sup>+</sup>CD4<sup>+</sup> (top row) and DAPI<sup>-</sup>CAR<sup>+</sup>CD8<sup>+</sup> (bottom row) cells over the same period. Each dot represents the mean of three biological replicates with error bars as SD.

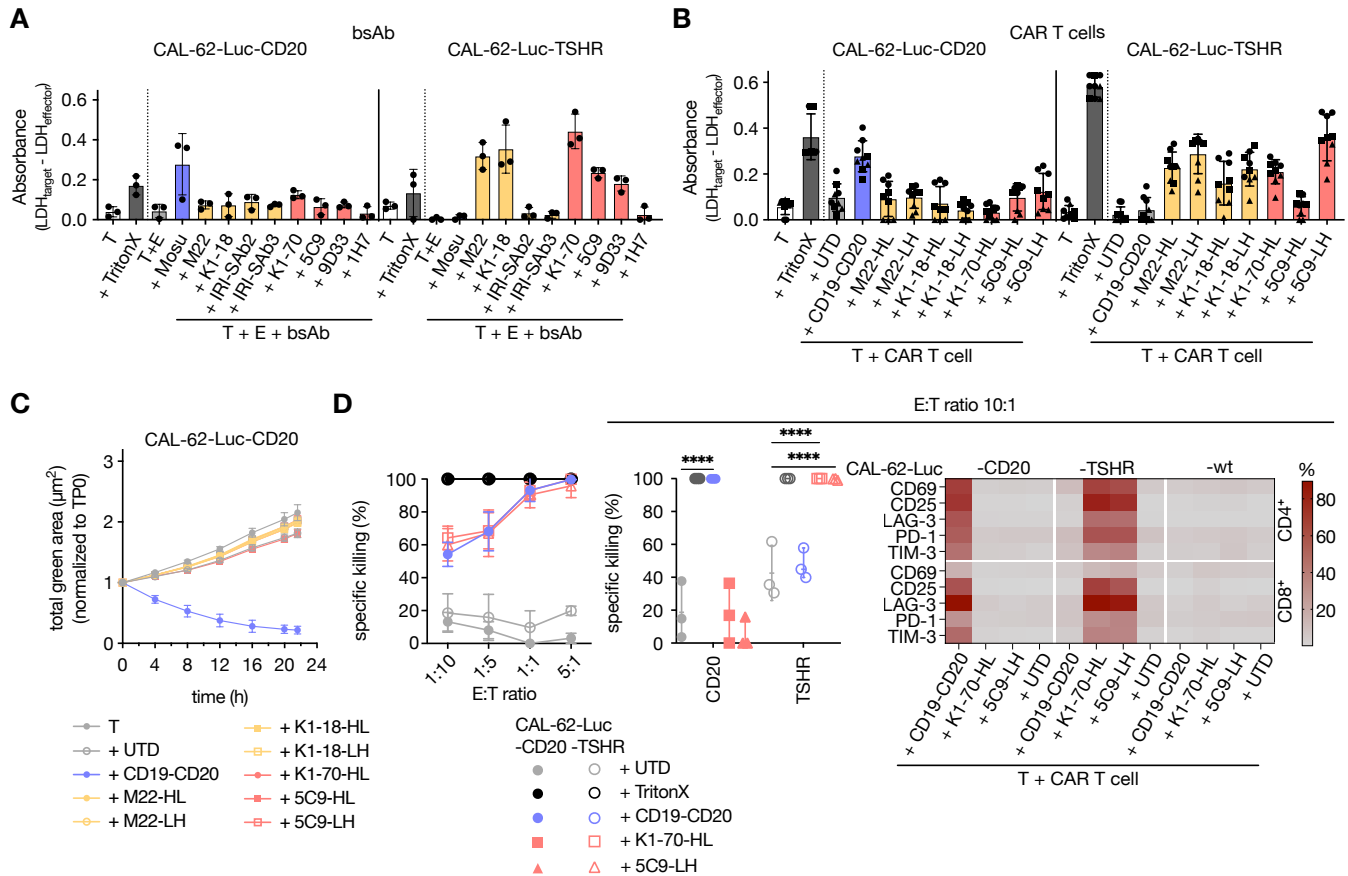

**Supplementary Figure 4. Cytotoxicity of anti-TSHR bispecific antibodies and CAR T cells.** **A** LDH release of lysed CAL-62-Luc-TSHR (right) and CAL-62-Luc-CD20 control (left) target cells after 48 h co-culture with the eight anti-TSHR bsAbs (10 nM) and HD CD3<sup>+</sup> T cells at an effector-to-target (E:T) ratio of 5:1. Absorbance was measured at 492 nm with absorbance values at 650 nm and effector LDH release subtracted. Mosunetuzumab (Mosu) served as a control. Each dot represents one technical replicate with error bars as SD. **B** The same read-out after 24 h co-culture with anti-TSHR CAR T cells in both scFv orientations (HL, LH). UTD and CD19-CD20-CAR T cells are the controls. Three biological replicates (●, ▲, ■) were used with three technical replicates each. These data accompany the luciferase-based killing shown in Figure 3B. **C** Proliferation of GFP<sup>+</sup> CAL-62-Luc-CD20 control target cells during co-culture with CAR T cells, monitored by live-cell imaging (Incucyte S3) over 24 h and expressed as total green area normalized to TP0. Each dot represents the mean of three biological and three technical (n = 9) replicates with error bars as SD. **D** Determination of the optimal E:T ratio for specific killing by HD-derived CAR T cells (n = 3) in 24-h co-cultures with CAL-62-Luc-TSHR or CAL-62-Luc-CD20 target cells, read out by luciferase assay. Used E:T ratios were 1:10, 1:5, 1:1 and 5:1 (left plot). Each dot represents the mean of three biological and two technical replicates (n = 6) with error bars as SD. Corresponding controls at an E:T ratio of 10:1 are shown in the middle plot (specific killing) and in the right plot (activation). Each dot represents one biological replicate and the mean of two technical replicates with error bars as SD. \*\*\*\*: p < 0.0001. E: effector cells, T: target cells.

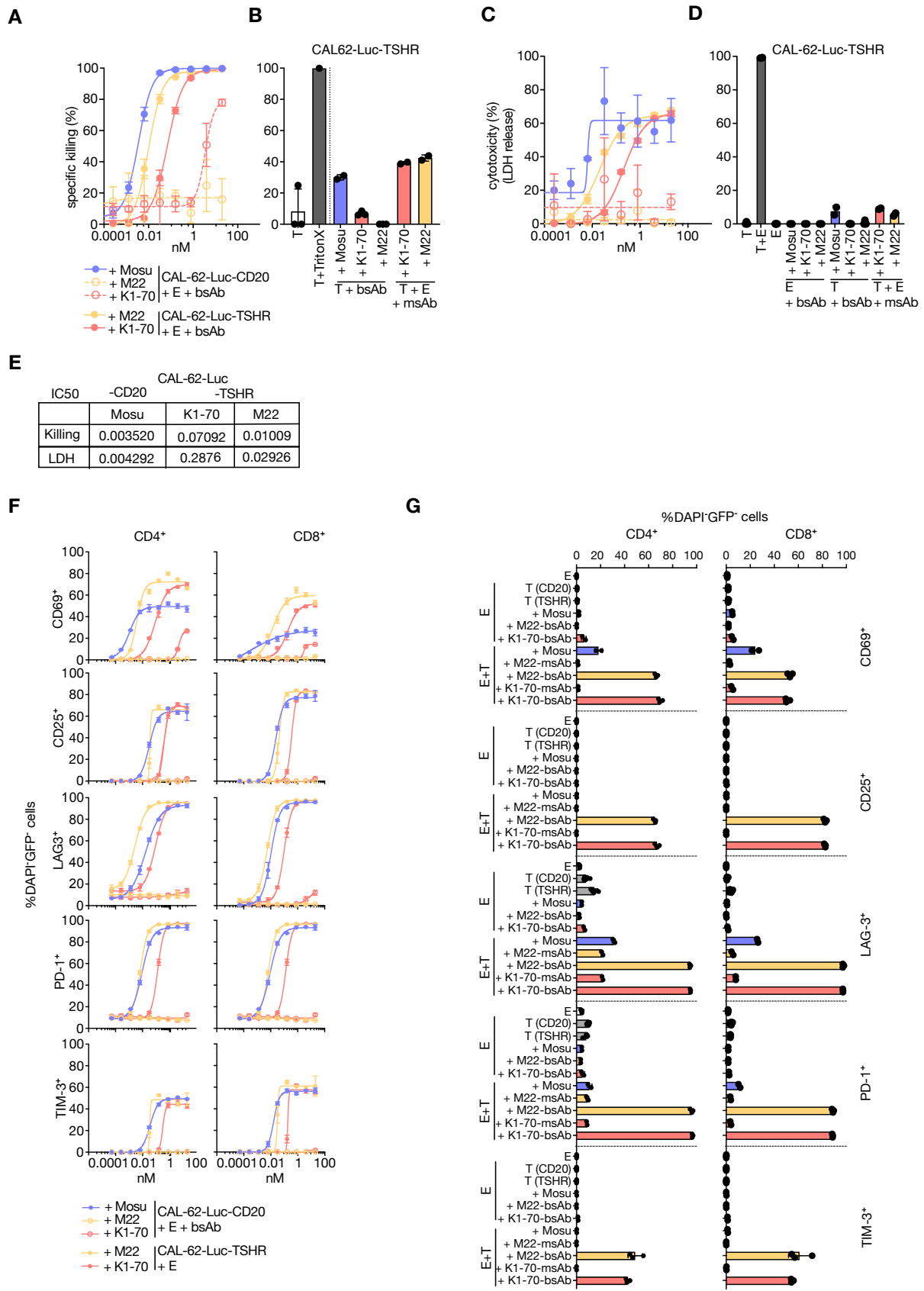

**Supplementary Figure 5. Titration of the anti-TSHR M22-bsAb and K1-70-bsAb.** **A** Specific killing of CAL-62-Luc-TSHR/CD20 cells in a 48 h co-culture with HD T cells in a 5:1 E:T ratio and a serial-dilution of M22- and K1-70-bsAbs starting at 20 nM. Percentage of specific killing was determined via luciferase assay. **B** Specific killing capacities of controls belonging to A. Bispecific and monospecific antibodies were used at the highest concentration of 20 nM in mono- (T) and co-culture (T+E). **C** Cytotoxicity (LDH release) of bsAbs co-cultured as described in A. **D** LDH release of controls described in B, with the addition of effector cells

only and effector cells in the presence of 20 nM of bsAbs. **E** The IC<sub>50</sub>-values for the bsAb in the presence of their corresponding specific target cells is shown for both, the luciferase assay and LDH release, as described in A and in C, respectively. **F** Expression of activation markers on CD4<sup>+</sup> and CD8<sup>+</sup> effector T cells upon bsAb titration and co-culture as described in A and determined by flow cytometry. **G** Controls belonging to F, as described in B and D. In A, C and F, linear regression analyses were performed and each dot represents the mean of three technical replicates with error bars as SD. In B, D and G, each dot represents one technical replicate (n = 3). E: effector cells, T: target cells.

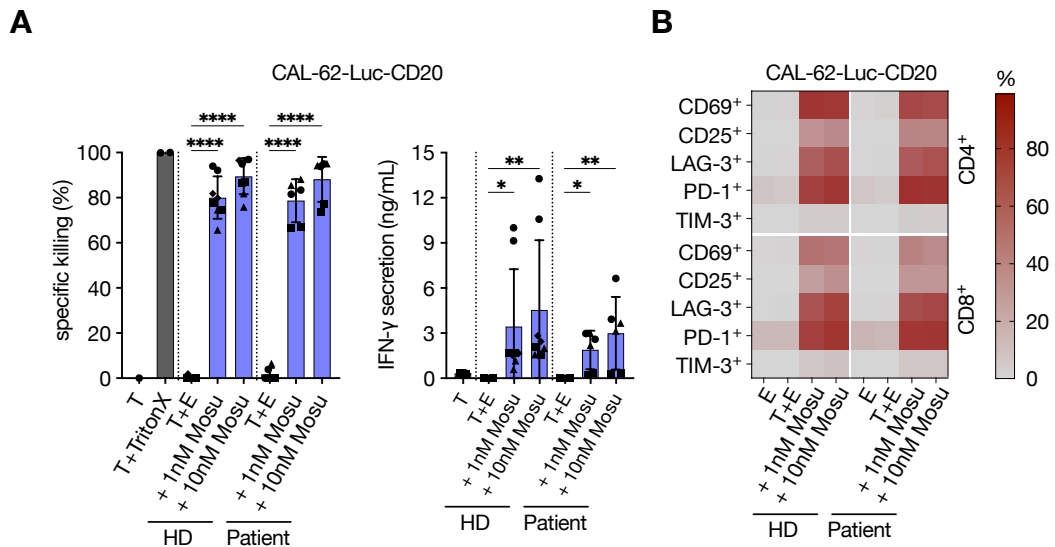

**Supplementary Figure 6. Co-culture experiment with bsAbs on the control cell line. A** Antigen dependent cytotoxicity of CAL-62-Luc-CD20 cells by 1 or 10 nM mosunetuzumab (Mosu) in 48 h co-culture with T cells from healthy donors (n = 4) or patients (each n = 3) as in Figure 5. Percentage of specific killing was determined by luciferase assay (left) and effector cytotoxicity read out by IFN-γ secretion (right). Per biological replicate indicated by different symbols (●, ▲, ■, ◆), two technical replicates were used with error bars as SD. For statistical analysis, ordinary two-way ANOVA with Dunnett's multiple comparisons test was used. **B** Effector activation marker readout as in Figure 5B. Each square represents the mean of four (HD) or three (patient) biological replicates with two technical replicates each. \*\*\*\*: $p < 0.0001$, \*\*: $p \leq 0.01$, \*: $p \leq 0.05$. T: target cells, E: effector cells.

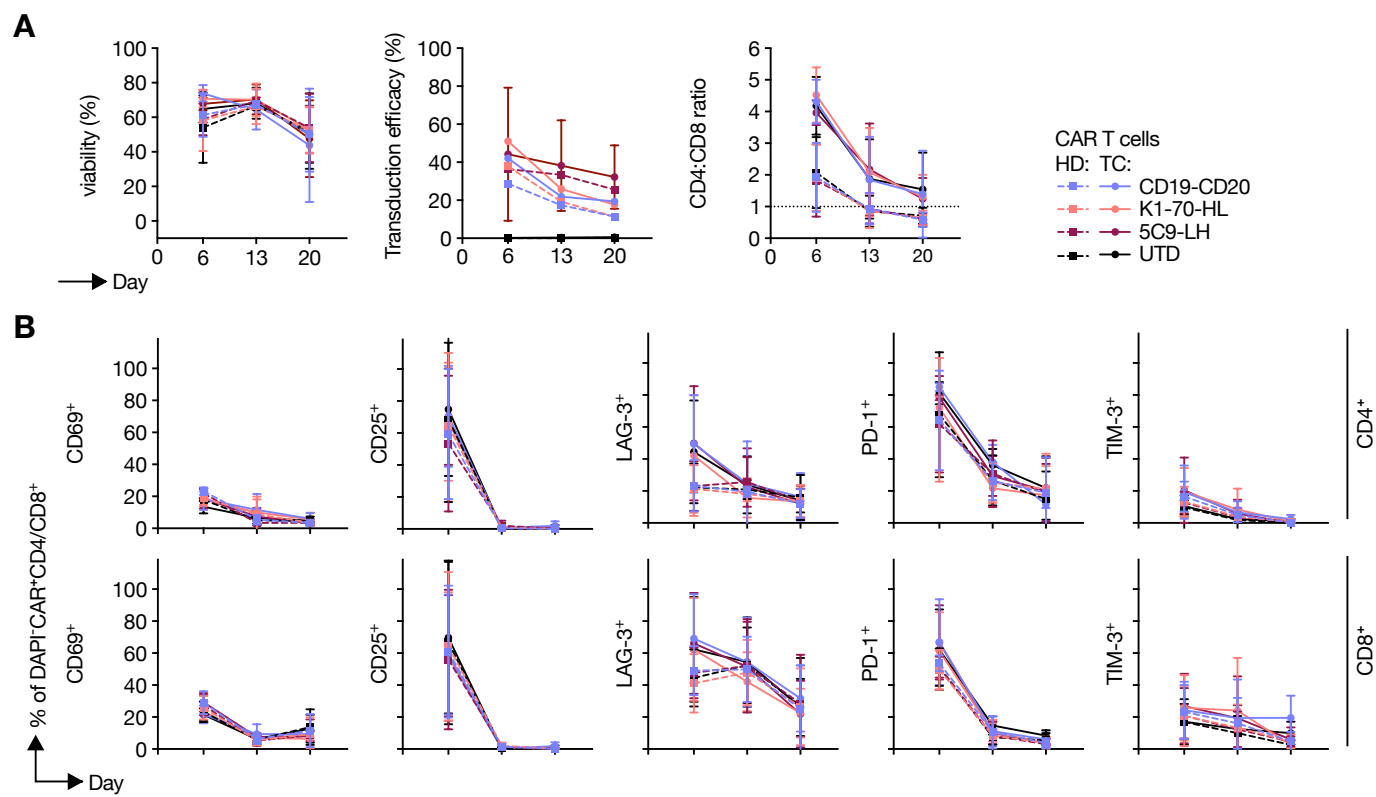

**Supplementary Figure 7. Production characteristics of patient- and HD-derived anti-TSHR CAR T cells.** **A** Percentage of viability and transduction efficiency and CD4:CD8 ratio of CAR T cells generated from HD or thyroid cancer patient (TC) T cells, determined by flow cytometry on days 6, 13 and 20 of expansion. **B** Activation markers (CD69, CD25, LAG-3, PD-1 and TIM-3) on DAPI<sup>-</sup>CAR<sup>+</sup>CD4<sup>+</sup> (top row) and DAPI<sup>-</sup>CAR<sup>+</sup>CD8<sup>+</sup> (bottom row) cells over the same period. Each dot represents the mean of three biological replicates with error as SD.

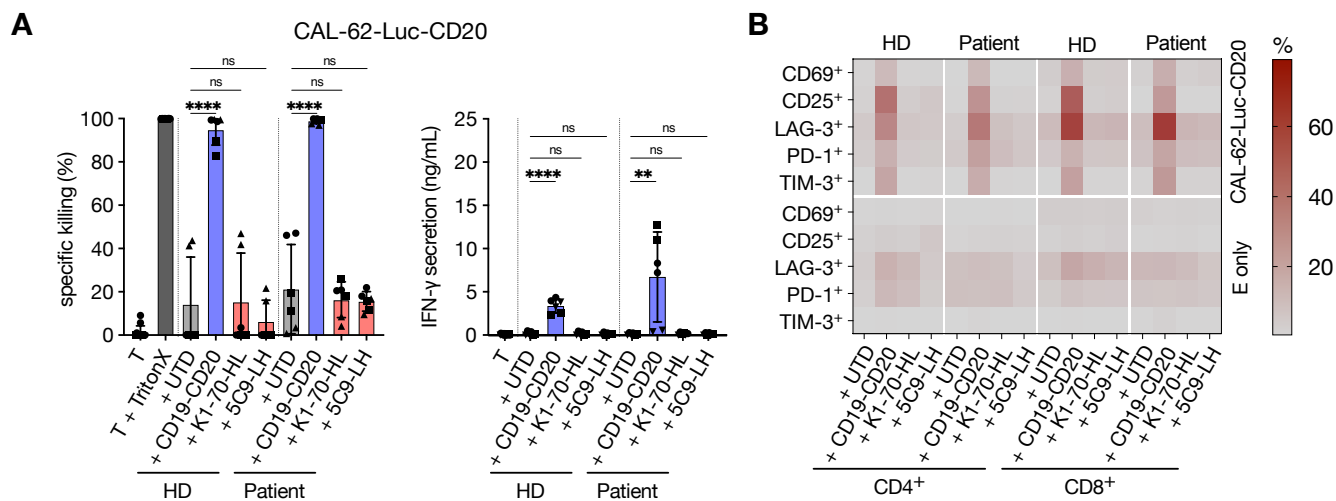

**Supplementary Figure 8. Anti-TSHR CAR T cell activity against the TSHR-negative control cell line. A** Specific killing and IFN-γ secretion in 24-h co-cultures of CAL-62-Luc-CD20 target cells with K1-70-HL-CAR, 5C9-LH-CAR, CD19-CD20-CAR or UTD T cells generated from HD or patients as in Figure 5C, D. Percentage of specific killing was determined by luciferase assay (left) and effector cytotoxicity read out by IFN-γ secretion (right). Per biological replicate indicated by different symbols ($\bullet$, ▲, ■), two technical replicates were used with error bars as SD. For statistical analysis, ordinary two-way ANOVA with Dunnett's multiple comparisons test was used. **B** Effector activation and exhaustion marker readout as in Figure 5D. Each square represents the mean of three biological replicates with two technical replicates each. \*\*\*\*: $p < 0.0001$, \*\*: $p \leq 0.01$, ns: non-significant. T: target cells, E: effector cells.

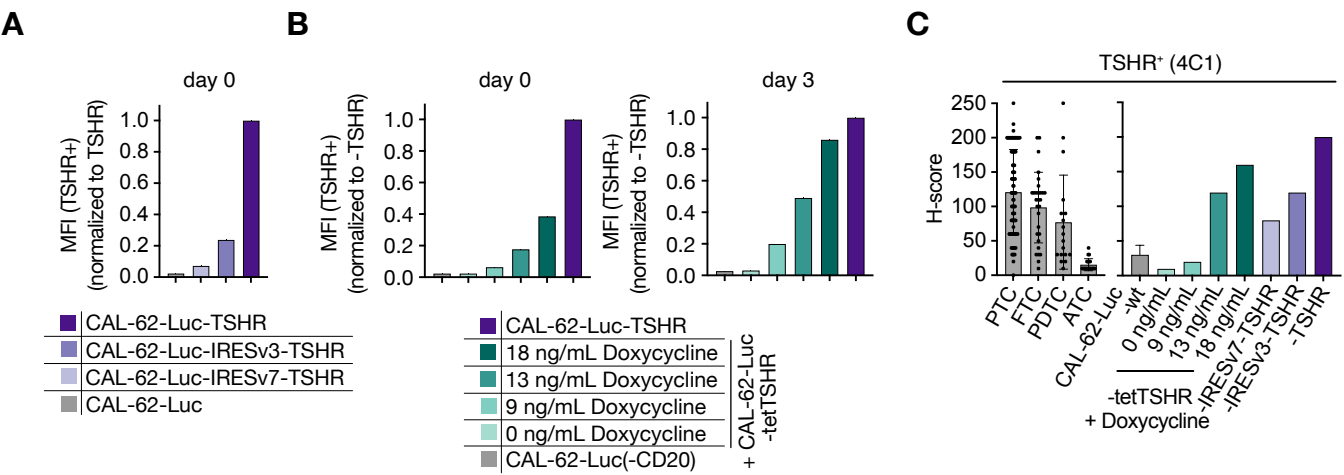

**Supplementary Figure 9. TSHR surface expression on the engineered CAL-62 target cell lines. A** Median fluorescence intensity (MFI) of surface TSHR (M22) on CAL-62-Luc-TSHR, CAL-62-Luc-IRESv3-TSHR, CAL-62-Luc-IRESv7-TSHR and parental CAL-62-Luc cells on day 0 of co-culture and normalized to CAL-62-Luc-TSHR. **B** Equivalent analysis of CAL-62-Luc-tetTSHR cells after a 72 h induction with 0, 9, 13 or 18 ng/mL doxycycline, on day 0 (left) and day 3 (right), with CAL-62-Luc-TSHR and CAL-62-Luc-CD20 shown as references. **C** H-score of TSHR immunohistochemistry stainings (4C1) across the thyroid cancer TMA cohort (left, same cohort as in Figure 1B) plotted on the same scale as the H-scores obtained for the engineered CAL-62 cell lines.

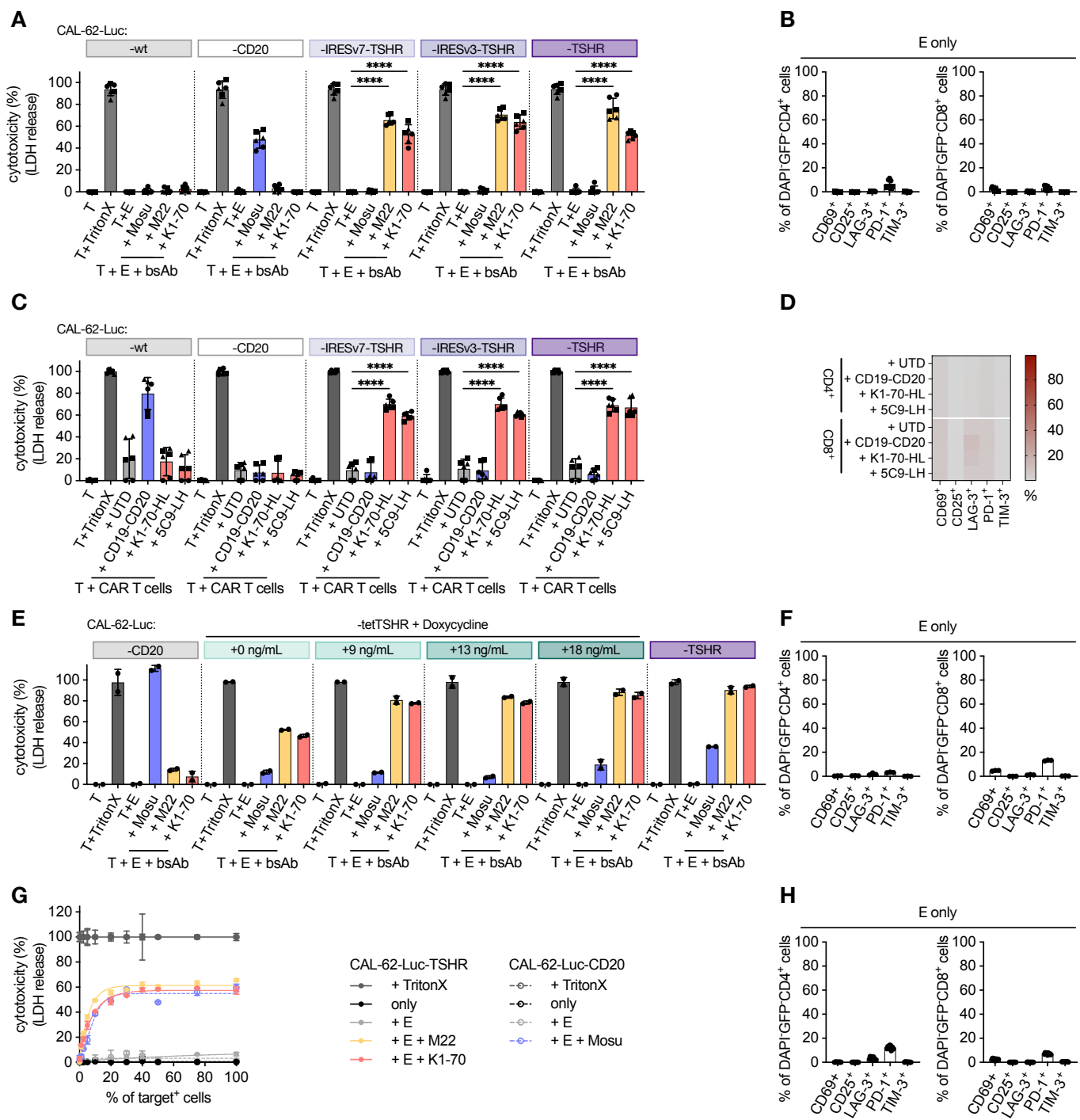

#### Supplementary Figure 10. Cytotoxicity of anti-TSHR bispecific antibodies and CAR T cells in varying TSHR expression cell models.

**A** Cytotoxicity against CAL-62-Luc-wt/-CD20/-IRESv7-TSHR/-IRESv3-TSHR/-TSHR target cells after 48 h co-culture with 1 nM of M22-bsAb or K1-70-bsAb and HD T cells at an E:T ratio of 5:1, based on LDH release. Mosunetuzumab (Mosu) served as a control. Three biological replicates (●, ▲, ■) were used with two technical replicates each. For statistical analysis, ordinary two-way ANOVA with Dunnett's multiple comparisons test was used with error bars as SD. **B** Activation marker expression of effector T cells used in A. **C** Cytotoxicity of CD19-CD20, K1-70-HL or 5C9-LH CAR T cells against CAL-62-Luc-wt/-CD20/-IRESv7-TSHR/-IRESv3-TSHR/-TSHR target cells after 24 h co-culture in an E:T ratio of 5:1, based on LDH release. A total of three biological replicates as indicated by the symbols (●, ▲, ■) were used with two replicates each. Ordinary two-way ANOVA with Dunnett's multiple comparisons test was used for statistical analysis with error bars as SD. **D** Activation marker expression of effector CAR T cells used in C. **E** Percentage of cytotoxicity against CAL-62-Luc-CD20/-TSHR or CAL-62-Luc-tetTSHR cells previously incubated for 72 h with 0, 9, 13 or 18 ng/mL doxycycline and co-cultured for 48 h with 1 nM of M22-bsAb or K1-70-bsAb and HD T cells at an E:T ratio of 5:1, based on LDH release. Each dot represents one technical replicate with error bars as SD. **F** Activation marker expression of effector T cells used in E. **G** Cytotoxicity measured via LDH release of M22-bsAb, K1-70-bsAb and mosunetuzumab against an increasing mix of CAL-62-Luc-TSHR or CAL-62-Luc-CD20 with CAL-62-Luc cells in a 48-h co-culture with HD T cells in a 5:1 E:T ratio. Each dot represents the mean of three technical replicates with error bars as SD. **H** Activation marker expression of effector T cells used in G. \*\*\*\*: $p < 0.0001$. T: target cells, E: effector cells.
